# Dual Control of LDL-cholesterol Levels by ANGPTL3 and ANGPTL8

**DOI:** 10.64898/2026.03.30.715445

**Authors:** Yanqing Xu, Fei Luo, Justin Fletcher, Melissa Inigo, Shawn Burgess, Guosheng Liang, Lisa N. Kinch, Julia Kozlitina, Jonathan C. Cohen, Helen H. Hobbs

**Affiliations:** Howard Hughes Medical Institute, University of Texas Southwestern Medical Center, Dallas, TX 75390; Department of Molecular Genetics, University of Texas Southwestern Medical Center, Dallas, TX 75390; Center for Human Nutrition, University of Texas Southwestern Medical Center, Dallas, TX 75390; Department of Molecular Biology, University of Texas Southwestern Medical Center, Dallas, TX 75390; Eugene McDermott Center for Human Growth and Development, University of Texas Southwestern Medical Center, Dallas, TX 75390

**Author notes:** Corresponding author: Helen Hobbs, Jonanthan Cohen, University of Texas Southwestern Medical Center (UTSW), Dallas, TX 75390-9046.

**Keywords:** Angiopoietin-like protein 3 (ANGPT3), Angiopoietin-like protein 8 (ANGPTL8), lipoprotein lipase (LPL), endothelial lipase (EL), triglyceride, LDL

## Abstract

**BACKGROUND:** Inactivation of ANGPTL3 (angiopoietin-like protein 3, A3) is a proven therapeutic strategy for lowering plasma lipid levels independently of the LDL receptor (LDLR), yet the optimal approach to inactivate A3 remains unclear. A3 is proteolytically cleaved and circulates as full-length (A3-FL), N-terminal (A3-Nter) and C-terminal (A3-Cter) fragments. The specific contribution of each form of A3, and of its paralog, ANGPTL8 (A8), in modulating circulating levels of ApoB-Containing Lipoproteins (ABCLs) remain poorly defined. Clarifying these relationships will inform next-generation A3-directed therapies.

**METHODS:** We performed liver perfusion studies to directly compare the number and composition of VLDL particles secreted from mice with and without A3. To amplify effects on cholesterol metabolism, we generated *Ldlr^-/-^* mice expressing wildtype A3 (A3-WT), A3-FL or A3-Nter, with or without co-expression of A8, and analyzed plasma lipids, circulating A3 and A8 complexes, and intravascular lipase activities. Complementary *in vitro* assays and structural modeling were used to assess relative endothelial lipase (EL) inhibition by A3 alone or in complex with A8.

**RESULTS:** Liver perfusion studies revealed that A3 inactivation does not alter the rates of hepatic secretion of VLDL in wildtype or *Ldlr^-/-^*mice. Inactivation of A8 alone lowered plasma LDL-cholesterol (C) levels by ∼20%, an effect dependent upon the expression of both EL and A3. Maximal inhibition of lipoprotein lipase (LPL) required co-expression of A8 plus both A3-FL and A3-Nter, indicating that A3 cleavage, in addition to A8 expression, is essential for maximal LPL inhibition. In contrast, A8 expression, but not A3 cleavage, was required for optimal EL inhibition.

**CONCLUSIONS:** A8 acts in concert with A3 to differentially modulate LPL- and EL-mediated lipolysis, which antagonizes hepatic clearance of newly-secreted atherogenic ABCLs. This mechanistic framework refines our understanding of A3-targeted lipid lowering and highlights the therapeutic potential of dual A3- plus A8-directed strategies to treat dyslipidemia and prevent atherosclerotic cardiovascular disease.

**Clinical perspective:** *What is new?:*

- Inactivation of A3 lowers circulating ABCL levels without altering hepatic secretion rates of VLDL-ApoB or -TG.
- Proteolytic cleavage of A3 is required for maximal inhibition of LPL.
- Inactivation of A8 lowers LDL-C levels through an A3- and EL-dependent, but LDLR-independent, mechanism.

*What are the clinical implications?:*

- Combining A8 inhibition with A3-inactivating therapies offers a strategy to achieve greater reduction in LDL-C levels and atherosclerotic cardiovascular risk.

## Background

The angiopoietin-like proteins (ANGPTLs) are a family of secreted proteins related to angiopoietins (ANGPTs). ANGPTL3 (A3) was first identified in 1999 and subsequently recognized as an endogenous inhibitor of lipoprotein lipase (LPL), the major intravascular triglyceride (TG) hydrolase.^1,2^ LPL is anchored to capillary endothelial surfaces by GPIHBP1 and heparan sulfate proteoglycans and catalyzes the rate-limiting step in delivery of TG-derived fatty acids to peripheral tissues.^3^ In addition to LPL, A3 inhibits endothelial lipase (EL), a phospholipase that remodels phospholipids in low density lipoprotein (LDL) and high density lipoprotein (HDL).^4–7^ Inactivation of A3 lowers plasma cholesterol levels through a mechanism that requires expression of EL.^8–10^

In mice, A3 inactivation increases both LPL and EL activity, lowers plasma lipid levels, and protects hyperlipidemic *Apoe^-/-^* mice from atherosclerosis.^11^ In humans, loss-of-function mutations in A3 are associated with marked lipid lowering and protection from atherosclerotic cardiovascular disease (ASCVD).^12,13^ Currently, the only approved A3-directed therapy is evinacumab, an anti-A3 monoclonal antibody indicated for the treatment of homozygous familial hypercholesterolemia to reduce ASCVD risk.^14^ An A3 antisense oligonucleotide (ASO) was developed to treat individuals with suboptimal responses to statins and other lipid-lowering agents, but was associated with hepatic steatosis and elevations in liver enzymes.^15^ More recently, A3-targeted small interfering (si) RNA therapies have shown improved hepatic safety profiles but achieve more modest LDL-cholesterol (C) lowering (∼17-40%) than observed in individuals with homozygous A3 deficiency.^16,17^ Collectively, these findings indicate that current pharmacological approaches fail to fully inactivate A3 and underscore the need for additional strategies to achieve the degree of LDL-C reduction seen with genetic A3 deficiency, and reach most recent guideline-recommended LDL-C targets.^18^

Multiple factors likely contribute to difficulties achieving sufficient A3 lowering for optimal lipid-lowering. First, the specific A3 species that inhibit intravascular lipases remain incompletely defined. Circulating A3 is synthesized in hepatocytes and cleaved at a canonical furin site to yield an N-terminal coiled-coil domain (A3-Nter) and a C-terminal fibrinogen-like domain (C-ter, FBG).^19,20^ Both full-length A3 (A3-FL) and A3-Nter contain a LPL-binding motif that is shared with ANGPTL4 (A4).^21^ The relative contributions of A3-FL and A3-Nter to LPL inhibition remain uncertain. Although A3-FL and wildtype A3 (A3-WT) show similar LPL inhibitory activities *in vitro*, expression of A3-Nter in *A3^-/-^* mice produces a larger increase in TG levels than equivalent expression of A3-FL,^19^ suggesting that A3-Nter may be the principal *in vivo* LPL inhibitor; however, these studies used supraphysiological levels of expression that may not reflect normal physiology.

Discrepancies between *in vitro* and *in vivo* A3 activity likely reflect the influence of ANGPTL8 (A8), an A3 paralog lacking the C-ter.^22–24^ Unlike A4, A3 requires A8 to achieve full inhibition of LPL.^25^ *In vitro*, the A3/A8 complex has an estimated 100-fold higher LPL inhibitory potency than A3 alone.^26^ A8 expression is strongly nutrition-dependent.^22–24^ In mice, A8 inactivation alters postprandial plasma TG levels but not plasma cholesterol levels, consistent with a selective role in A3-mediated LPL inhibition.^27,28^ In cynomolgus monkeys, A8 inactivation using an anti-A8 antibody reduced plasma TG levels by 65% but has no effect on plasma levels of LDL-C.^28^ Similarly, humans who are heterozygous for A8 loss-of-function mutations have a 19-24% reduction in TG levels.^29^ They also have a 4-17% reduction in LDL-C levels, suggesting that A8 deficiency has a broader effects on lipoprotein metabolism. The reduction in LDL-C raises the question of whether, and in what form, A8 participates in EL inhibition. *In vitro* studies report conflicting rank orders for EL inhibition by A3-FL, A3-Nter and A3-WT in the presence or absence of A8. One study using conditioned media found that A3-Nter inhibited EL more potently than A3-FL, but less well than A3-WT, and that A8 did not contribute to this enzymatic inhibition.^30^ Another study found that A3-FL was the more effective EL inhibitor.^31^

Another challenge for A3-directed therapy is the magnitude of A3 suppression required to meaningfully lower LDL-C levels. Humans heterozygous for A3 loss-of-function mutations do not always have reduced plasma LDL-C levels.^12,32^ Thus, a 50% reduction in A3 is insufficient to achieve meaningful cholesterol lowering. Estimates suggest that A3 levels must be reduced by over 75-90% to significantly lower plasma LDL-C,^15,32^ a degree of suppression that has been difficult to achieve safely. A practical consideration is that A3 is a abundant protein in plasma, with circulating levels ranging from 3-20 nM, exceeding those of A8 (0.3-1 nM).^33,34,35^ Only ∼10% of A3 is bound by A8,^26,33^ whereas more than half of A8 is estimated to be complexed with A3.^34^ The anti-A3 antibody evinacumab must be administered intravenously to achieve adequate A3 inactivation.

The mechanism by which A3 inactivation lowers plasma ABCLs remains incompletely defined. In mice and humans, LDL-C lowering with A3 inhibition occurs independently of the hepatic LDLR and other canonical lipoprotein receptors, including LRP1 (LDL receptor related protein 1), scavenger receptor SCARB1 and syndecan 1 and is preserved in *Apoe^-/-^* mice.^9,21,36^ To test if reduced hepatic secretion of VLDL lipids contributes to the lipid-lowering effects of A3 inactivation, we performed liver perfusion studies to capture, characterize and compare newly-secreted VLDL from *Ldlr^-/-^*;*A3^-/-^*and *Ldlr^-/-^* mice. We then used genetically modified *Ldlr^-/-^*mice expressing either A3-FL or A3-Nter, with or without A8 co-expression, to define their relative contributions to TG and cholesterol metabolism. These findings provide new mechanistic insights with translational implications for the design of next-generation lipid-lowering therapies targeting A3 and A8.

## MATERIAL AND METHODS

Detailed methods for liver perfusion, VLDL isolation, immunoblotting, ELISA, lipid and lipase activity assays, quantitative real-time PCR, AAV infection in mice, cell lines, transient transfections, and structure modeling are provided in Supplementary Material.

The datasets generated and/or analyzed during the current study are derived from mouse experiments and are available from the corresponding author on reasonable request for non-commercial scientific research.

### Mice

Mice were housed and fed standard chow diets *ad libitum.* All animal experiments described in this study have been approved and conducted under the oversight of the UT Southwestern Institutional Animal Care and Use Committee.

The *Ldlr^-/-^* mice, *Lipg^-/-^* mice, and *A3^-/-^* mice were generated as described.^36,37,38^ All mice were on a C57BL/6J genetic background. Using CRISPR-Cas9, we generated *Ldlr^-/-^;A3^FL/FL^* and *Ldlr^-/-^;A3^Nter/Nter^*mice as described in the legend to Figure 2. A8-deficient *Ldlr^-/-^* mice were generated in house and then backcrossed for ≥3 generations before crossing with the indicated lines to generate the mice used in the studies. Breeding schemes and genotyping methods are provided in Supplementary Methods.

We did not observe any baseline differences between male and female mice under similar experimental conditions. Therefore, experiments were conducted only in male mice.

### Statistical Analysis

Data are presented as mean ± SD. Two-tailed parametric t tests and one-way ANOVA were used when normality assumptions were met, as determined by Shapiro-Wilk test and graphical diagnostic methods, using GraphPad Prism 10. For non-normally distributed data, or when normality could not be reliably assessed because of small sample size, nonparametric tests (Mann-Whitney U-test or Kruskal-Wallis test) were used to compare groups.

No experiment-wide or manuscript-wide multiple testing correction was applied; within-experiment *p* values were adjusted for multiple comparisons using Dunnett’s or Tukey’s tests as indicated. For experiments involving only prespecified pairwise comparisons, Fisher’s least significant difference (LSD) test was used only when the overall one-way ANOVA was significant (*p*<0.05). Statistical details for each experiment are provided in the figure legends and Supplementary Table 6.

## RESULTS

### No Reduction in VLDL-ApoB or -TG Secretion with A3 Inactivation

Previously, we found no evidence that A3 inactivation reduces the rate of hepatic ApoB secretion, yet we also found that the rate of TG accumulation in plasma was significantly reduced after we inhibited intravascular lipases.^36^ To resolve this apparent paradox, we directly quantified VLDL-ApoB and VLDL-TG secretion using *ex vivo* liver perfusion studies. Age- and sex-matched chow-fed WT and *A3^-/-^* male mice were anesthetized and treated with heparin before liver isolation and *ex vivo* perfusion. Perfusates were collected every 15 min and VLDL-ApoB-100, -ApoB-48, and -ApoE were quantified (Figure 1A). Across strains, secretion of these three proteins did not differ. In one pair, ApoE was minimally detectable for unknown reasons, but the result did not alter overall conclusions. The ratios of TG and cholesterol to ApoB-100 and to ApoB-48 were also similar in *A3^-/-^* mice and WT mice (Figure 1B).

**Figure 1.**
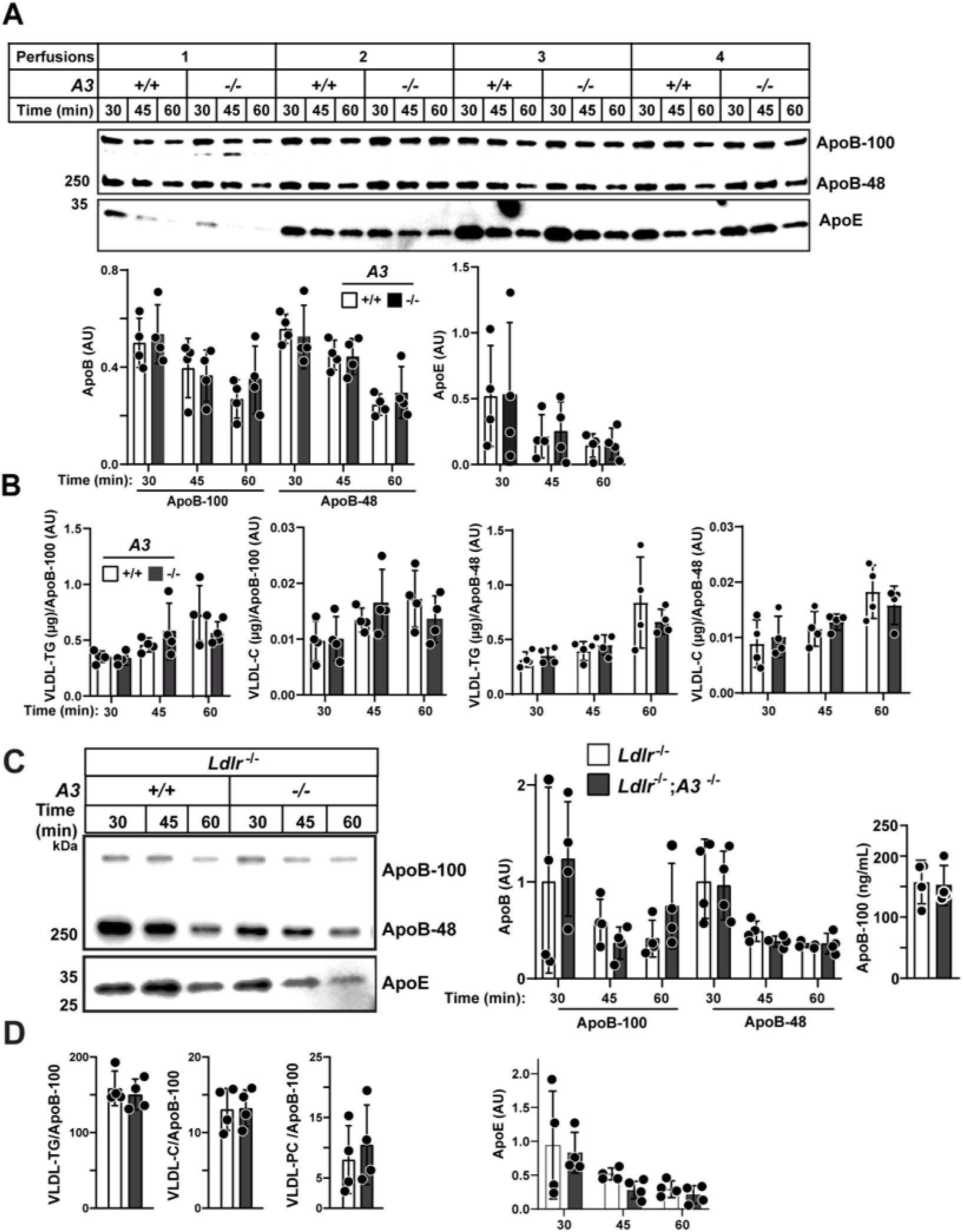
No changes in the rate of secretion of VLDL-ApoB or VLDL-TG with inactivation of A3. Liver perfusion and immunoblotting were performed as described in Supplementary Methods. Perfusates were sampled at 30, 45, and 60 min. At each point, VLDL was isolated and ApoB and ApoE proteins were quantified using immunoblot analysis and LI-COR imaging. **A**, Immunoblot analysis of VLDL-ApoB and -ApoE from liver perfusates of chow-fed male *A3*^-/-^ and WT controls (n=4/group, 11-13 wk). **B,** VLDL lipids from the liver perfusates were measured enzymatically in WT and *A3^-/-^* mice and normalized to ApoB levels. **C,** VLDL-ApoB and -ApoE were measured in *Ldlr^-/-^;A3*^-/-^ and *Ldlr^-/-^* controls (n=4/group, age matched 8-19 wks) after 2 wks on a high-sucrose diet. ApoB-100 levels were measured by LICOR (values are normalized to the +/+ group at 30 min) and by ELISA. **D,** VLDL lipids were measured in *Ldlr^-/-^;A3*^-/-^ and *Ldlr^-/-^*controls. Differences between WT and KO (n=4 per group) at each timepoint were tested using the Mann-Whitney test (with a Bonferroni correction for multiple time-point comparisons) No significant differences were observed (*p*>0.05 for all comparisons). Statistical values were not added to the graphs when no significant differences were found. When differences were statistically significant, the exact *p* values are provided in Supplementary Table 6. Each liver perfusion experiment included one mouse of each genotype. Data from four independent experiments were processed and analyzed in parallel.

To exclude any effects of heparin, we repeated the perfusion without heparin and obtained similar results (Figure S1A). VLDL-ApoB-100 levels were measured by ELISA (Enzyme Linked Immunoabsorbent Assay) in pooled samples from each mouse. The TG and cholesterol to ApoB-100 ratios in VLDL did not differ between strains (Figure S1B), indicating that heparin did not confound the measurements. Heparin was therefore omitted from subsequent studies.

Since the LDL-C-lowering effect of A3 inactivation is independent of LDLR,^9,21,36^ we inactivated A3 in *Ldlr^-/-^*mice to accentuate the effects on ABCL levels. To stimulate VLDL secretion, *Ldlr^-/-^* and *Ldlr^-/-^*;*A3^-/-^* mice were fed a high-sucrose diet (HSD) for two wks.^39^ Secretion of VLDL-ApoB and -ApoE was indistinguishable between genotypes (Figure 1C, left, and S1C). Likewise, ApoB-100 levels in pooled samples were comparable across groups (Figure 1C, right) and the ratios of TG, cholesterol and phosphatidylcholine (PC) to ApoB-100 did not differ between strains (Figure 1D).

From these experiments, we conclude that the reduced plasma lipid levels in *Ldlr^-/-^*;*A3^-/-^* mice are not attributable to decreased hepatic secretion of VLDL particles or diminished lipid content per particle. Next, we asked which circulating A3 species are required for the lipid-lowering effect of A3 deficiency.

### Circulating Levels of A3 Are Altered in Mice Expressing Only A3-FL or Only A3-Nter

A3 is cleaved at one of two adjacent furin cleavage sites: between residue Arg^221^ and Ala^222^ or Arg^224^ and Thr^225^ (Figure 2A).^19^ To define how cleavage relates to function, we generated *Ldlr^-/-^* mice expressing only full-length A3 (A3-FL) by substituting alanine for both arginines (R221A, R224A) (*Ldlr^-/-^;A3^FL/FL^* mice) (Figure 2B) and mice expressing only A3-Nter (residues 1-224) by introducing a premature stop codon at position 225 (ACT → TGA) (*Ldlr^-/-^;A3^Nter/Nter^*mice). Mutations were confirmed by sequencing. Three founder lines were established per genotype, and one line was selected for detailed study. Both new strains were healthy with normal appearance, fertility, Mendelian inheritance, body and liver weights compared to age- and sex-matched *Ldlr^-/-^* mice (Tables S2 and S3, Figure S2A). No gross differences were appreciated in morphology or routine histology of liver, heart or kidney (n=5-6/group, 11-16-wk-old male mice) compared to *Ldlr^-/-^* controls (data available upon request).

**Figure 2.**
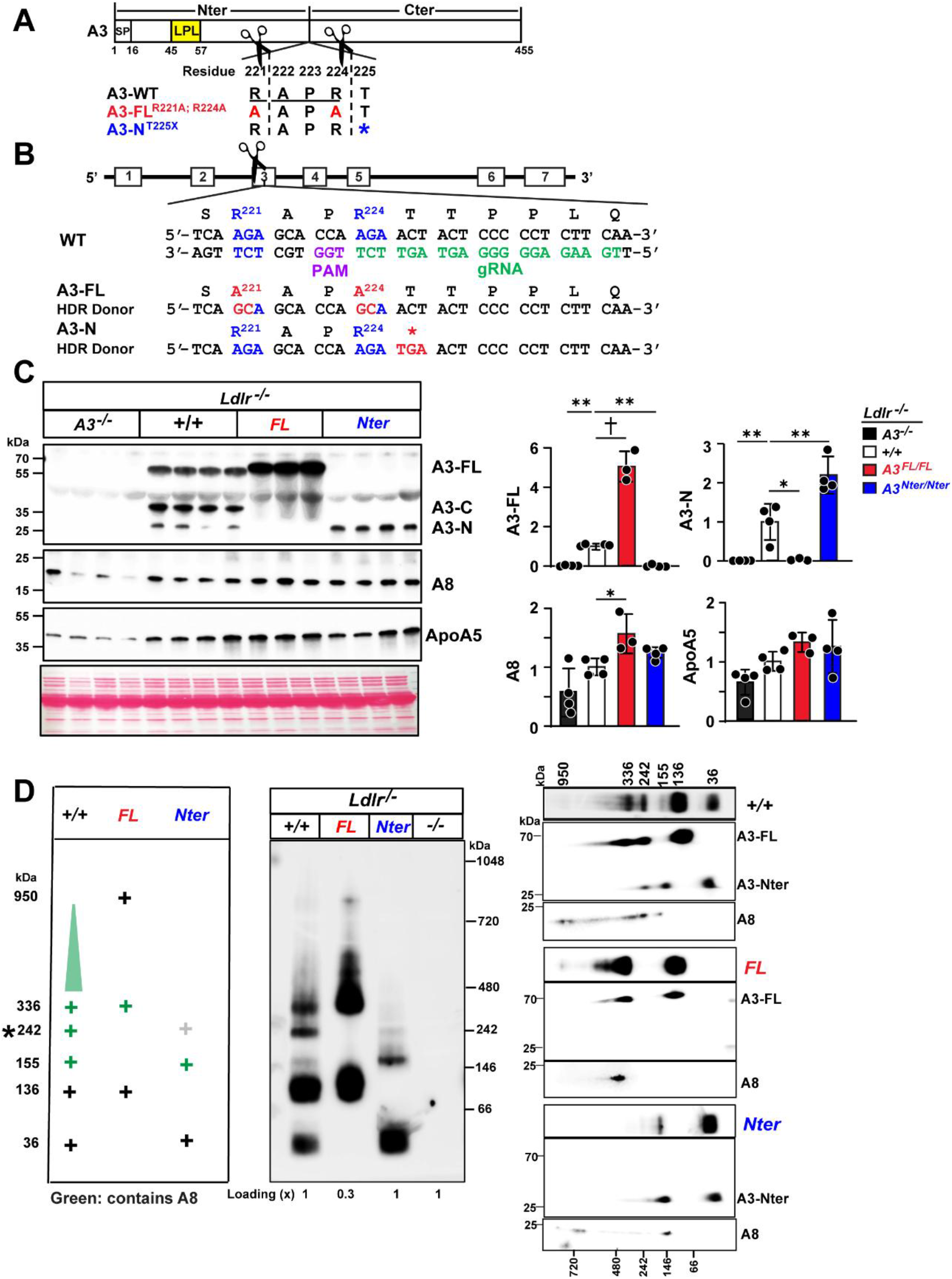
A3 cleavage alters circulating levels of A3 isoforms. **A,** Schematic of A3 variants expressed in mice: A3-WT, cleavage resistant A3-FL and truncated A3-Nter. SP, signal peptide. LPL, lipoprotein lipase. **B,** Generation of *A3^FL/FL^* and *A3^Nter/Nter^* mice using CRISPR-Cas9. Single guide sgRNA (gRNA, green) was co-injected with single-stranded oligonucleotide (ssODN) A3-FL or A3-N donors to introduce the indicated mutations (highlighted in red) via homology-directed repair (HDR). The protospacer adjacent motif sequence (PAM, purple) corresponding to the gRNA target site is also highlighted. **C,** Immunoblotting of circulating A3, A8 and ApoA5 protein levels. A3 was detected by an anti-A3 pAb raised against full-length A3. Band intensities were quantified using LI-COR. Values are normalized to +/+ group (male, n=4, 4, 3, and 4/group, 8-12 wks, refed 4 h). **D,** Size distribution of A3-containing complex(s) by Native Page. The native lane was then exercised and resolved by denaturing SDS-PAGE in a second dimension (See Supplementary Methods). The schematic (left) indicates complex in A3-WT (+/+), A3-FL (FL) and A3-Nter (Nter) mice (green denotes A8-containing complexes). Mean values were compared using one-way ANOVA with Dunnett’s multiple comparisons test as implemented in GraphPad Prism. Data are presented as mean ± SD. \**p*<0.05, \*\**p*<0.01, †*p*<0.0001. Statistical values were not added to the graphs when no significant differences were found. When differences were statistically significant, the exact *p* values are provided in Supplementary Table 6. This experiment was repeated 3 times with similar results.

Hepatic A3 mRNA levels were similar in *Ldlr^-/-^*;*A3^FL/FL^* and *Ldlr^-/-^* controls (Figure S2B). However, the levels of total and mature mRNA but not pre-mRNA, were significantly reduced (−36% total mRNA, *p=*3.15E-07 and −53% mature mRNA, *p*=1.68E-09) in *Ldlr^-/-^*;*A3^Nter/Nter^* mice relative to *Ldlr^-/-^*controls, likely due to mRNA decay from the introduction of a premature stop codon. Immunoblotting analysis of plasma with a polyclonal antibody (AF136) revealed three bands at the expected sizes of A3-FL (65 kDa), A3-Nter (27 kDa), and A3-Cter (38 kDa) (Figure 2C); as expected, A3-Nter and A3-Cter were absent in *Ldlr^-/-^*;*A3^FL/FL^*mice. Despite comparable hepatic A3 mRNA levels, plasma A3-FL levels were 5-fold higher in *Ldlr^-/-^*;*A3^FL/FL^* mice than in *Ldlr^-/-^* mice (Figure 2C). Plasma A3-Nter levels were 20% higher in *Ldlr^-/-^*;*A3^Nter/Nter^* mice than in *Ldlr^-/-^* mice (Figure 2C), despite a reduction in hepatic A3 mature mRNA. The levels of total A3 (A3-FL+A3-Nter) in mice expressing A3-FL and A3-Nter were 3.8-fold higher (*p*=2.53E-08) and 58% lower (marginally significant, *p*=0.049) than mice expressing A3-WT (Figure S2C). To compare the levels and molarity of circulating A3, we assayed plasma A3 levels in the 3 mouse strains using different assays: densitometry of immunoblots with an A3-Nter mAb (Figure S2D), ELISA (Figure S2E) and SRM-MS (selected reaction monitoring mass spectrometry, Figure S2F). Despite assay-specific technical limitations that affected absolute quantification of individual A3 isoforms, the mass/molarity of total A3 in our mouse expressing A3-FL and A3-Nter were consistently higher or similar to that observed in A3-WT mice.

Plasma levels of A8 were 50% higher in *Ldlr^-/-^*;*A3^FL/FL^* mice than in *Ldlr^-/-^* mice but did not differ between A3-Nter and A3-WT expressing animals. Plasma levels of ApoA5, which binds A3/A8 and interferes with its LPL inhibitory effect,^40^ were similar among the strains (Figure 2C).

### Multiple Complexes of A3 Circulate in Plasma

Having confirmed that the genetically-modified mice expressed the expected A3 isoforms, we next compared the size distribution and A8 content of circulating A3-FL and A3-Nter complexes. Plasma was fractionated by size-exclusion chromatography, and A3-containing fractions were pooled (5-11, Figure S3A), subjected to non-denaturing gel electrophoresis and then immunoblotted for A3 (Figure 2D, middle). In *Ldlr^-/-^* mice, a series of A3-immunoreactive were seen with apparent molecular masses of 336, 242, 155, 136, and 36 kDa. Two of these bands (336 kDa and 136 kDa) were also present in *Ldlr^-/-^*;*A3^FL/FL^* mice with the 336 kDa band extending as a smear up to ∼950 kDa. The 242, 155, and 36 kDa bands, absent in *Ldlr^-/-^*;*A3^FL/FL^* mice, were present in *Ldlr^-/-^*;*A3^Nter/Nter^*mice. Thus, no A3-immunoreactive bands were shared between mice expressing only A3-FL and only A3-Nter.

To define the A3 and A8 content of these complexes, we excised each lane from the nondenaturing gel and subjected the proteins to SDS-PAGE in a second dimension (Figure 2D, right). The 336 kDa band in the sample from *Ldlr^-/-^* and *Ldlr^-/-^*;*A3^FL/FL^* mice contained both A3-FL and A8, whereas the 242 kDa band in *Ldlr^-/-^* mice contained A3-FL, A3-Nter, and A8. A faint band of similar size was seen in *Ldlr^-/-^*;*A3^Nter/Nter^* mice (Figure 2D, middle) but was below the detection limits after SDS-PAGE (Figure 2D, right). The 155 kDa band contained both A3-Nter and A8 and the 136 kDa band and 36 kDa band contained only A3-FL or A3-Nter, respectively.

Taken together, these findings indicate that both A3-FL and A3-Nter circulate in multiple distinct complexes, some incorporating A8, and that only the 242 kDa complex in *Ldlr^-/-^* mice contains A3-FL, A3-Nter and A8 together. Attempts to determine the precise stoichiometry of these complexes by mass spectrometry were inconclusive owing to limited protein yield.

To assess the physiological relevance of A3 cleavage, we examined how preventing or enforcing A3 cleavage affected plasma lipid levels.

### A3-FL and A3-Nter Are Both Required for Maximal Effect of A3 on Plasma TG

Plasma TG levels in *Ldlr^-/^*^-^*;A3^FL/FL^* and *Ldlr^-/-^;A3^Nter/Nter^* mice were intermediate between those in *Ldlr^-/-^* and *Ldlr^-/-^*;*A3^-/-^* animals (*p*=6.40E-11) (Figure 3A). TG levels were reduced by 50% and 28%, respectively in *Ldlr^-/-^;A3^FL/FL^* and *Ldlr^-/^*^-^*;A3^Nter/Nter^*mice, relative to *Ldlr^-/-^* controls. These data indicate that neither A3-FL alone nor A3-Nter can achieve the degree of LPL inhibition observed when both isoforms are present. To test this idea, we generated mice expressing different combinations of A3 isoforms on an *Ldlr^-/-^;A3^-/-^* background and assayed A3 levels by immunoblot (Figure S3B). The pattern of relative levels of A3 in the mice expressing A3-WT, A3-FL, and A3-Nter were similar to what was seen above in Figure 2C. In the hemizygous mice carrying a single allele expressing A3-FL (*A3^FL/-^*) or a single allele expressing A3-Nter (*A3^Nter/-^*), A3 levels were reduced by ∼50% compared to *A3^FL/FL^* and A3*^Nter/Nter^* mice. The reduction in A3-Nter was restored when A3-FL was co-expressed with A3-Nter in *A3^FL/Nter^* mice. *A3^FL/-^* and *A3^Nter/-^* mice had plasma TG levels that were ∼50% of their homozygous counterparts, consistent with a gene dosage effect. Notably, mice expressing both A3-FL and A3-Nter (FL/Nter) had plasma TG levels indistinguishable from mice expressing A3-WT, indicating that optimal LPL inhibition *in vivo* requires co-expression of A3-FL and A3-Nter.

**Figure 3.**
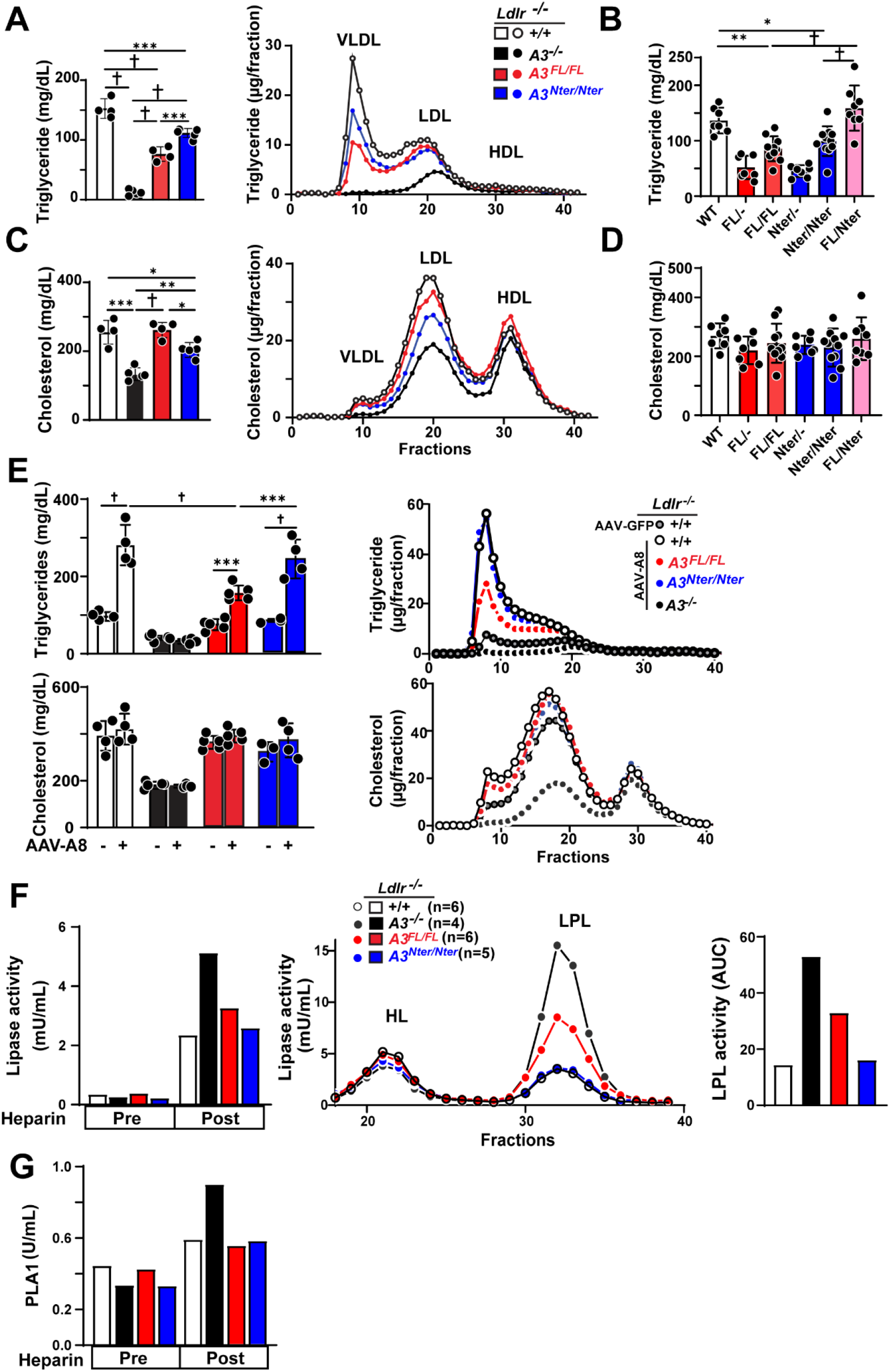
A3 cleavage has different effects on plasma triglyceride and cholesterol levels. **A**, Levels (left panel) and size distributions (right panel) of TG in plasma from *Ldlr*^-/-^ (+/+, A3-WT), *Ldlr*^-/-^*;A3^FL/FL^*, *Ldlr*^-/-^*;A3^Nter/Nter^* and *Ldlr*^-/-^*;A3*^-/-^ mice (male, n=4, 5, 4 and 5/group, 8-12 wks, refed 4 h). **B**, Plasma triglyceride levels in mice with different combinations of A3 isoforms *A3^+/+^(WT), A3^FL/-^, A3^FL/FL^, A3^Nter/-^, A3^Nter/Nter^, A3^FL/Nter^* (male, n=7, 7, 10, 7, 11, 8/group pooled from two experiments, 6-11 wks, refed 4 h). **C,** Levels (left panel) and size distributions (right panel) of cholesterol in 4 strains of mice. **D**, Plasma cholesterol levels in mice expressing different A3 isoforms. **E,** Hepatic overexpression of mouse A8 in 4 strains of mice. Plasma TG and cholesterol levels and their size distributions were measured after a 16 h fast to avoid the endogenous A8 expression (male, n=4, 4, 5, 4, 5, 5, 3, and 4/group, 8-12 wks). **F,** Plasma LPL activity in 4 mouse strains (male, n=6, 4, 6, 5/group, 8-12 wks, refed 4 h). Pre- and post-heparin blood was collected, and total triglyceride lipase activity was measured enzymatically as detailed in Supplementary methods (left). Post-heparin plasma was fractionated on heparin-Sepharose affinity chromatography, and HL and LPL activities were measured in the eluate; LPL activity was quantified as area under the curve AUC (right) by GraphPad. **G,** Phospholipase A1 activity was determined in mouse plasma samples from 4 strains of mice as described in the Supplementary Methods. Mean values were compared using ordinary one-way ANOVA with Tukey’s multiple-comparisons test (A and C) or one way ANOVA-Fisher’s LSD test (B-E) as implemented in GraphPad Prism. Data are presented as mean ± SD. \**p*<0.05, \*\**p*<0.01, \*\*\**p*<0.001, †*p*<0.0001. Statistical values were not added to the graphs when no significant differences were found. When differences were statistically significant, the exact *p* values are provided in Supplementary Table 6. All experiments were repeated 3 times with similar results.

Differences in plasma TG levels across strains were not accompanied by changes in hepatic TG (or cholesterol) levels (Figure S3C), in line with our finding that A3 deficiency does not interfere with the rate of hepatic VLDL-TG secretion (Figure 1).

### A3 Cleavage Is not Required for Its Cholesterol-Elevating Effect

Plasma cholesterol levels in mice expressing A3-FL were similar to those expressing A3-WT mice, whereas A3-Nter mice showed a modest, but significant, 18% reduction in plasma cholesterol levels (*p*=0.024), compared to the 48.6% reduction in *Ldlr^-/-^;A3^-/-^* mice (Figure 3C); the largest strain-dependent differences were in the LDL fraction (Figure 3C, right). Plasma cholesterol did not differ significantly between heterozygous or homozygous carriers of either A3-FL or A3-Nter (Figure 3D), indicating that A3 cleavage exerts little effect on cholesterol levels.

To test if A8 is rate-limiting under these conditions, we overexpressed A8 in the 4 strains of mice using an adeno-associated virus (AAV), achieving comparable levels of hepatic A8 mRNA and protein (Figure S3D). As expected, A8 expression in *Ldlr^-/-^* mice produced an increase in plasma TG, whereas no TG increase was apparent in *Ldlr^-/-^;A3^-/-^*mice, consistent with the requirement of A3 in A8-dependent LPL inhibition^22^ (Figure 3E, left). Despite markedly higher circulating A3 levels in *Ldlr^-/-^*;*A3^FL/FL^*mice (Figure 2C), A8 overexpression produced a more modest increase in plasma TG than in *Ldlr^-/-^*;*A3^Nter/Nter^*mice (157.24 ± 17.19 mg/dL vs 239.84 ± 43.60 mg/dL). A8 overexpression did not cause a significant difference in plasma TG levels in *Ldlr^-/-^*;*A3^Nter/Nter^* mice and *Ldlr^-/-^* mice (Figure 3E, top). The attenuated TG-raising effect of A8 expression in A3-FL mice is likely because most of the A3-FL are in higher molecular complexes that poorly incorporate A8 (Figure 2D). Alternatively, the A3-FL/A8 complexes may be less potent LPL inhibitors. In contrast, A8 overexpression has little or no effect on plasma cholesterol levels across genotypes (Figure 3E, bottom).

Together, these data indicate that A3 cleavage state and its interaction with A8 strongly modulate the efficiency of LPL inhibition and thereby plasma TG levels, whereas plasma cholesterol levels are comparatively insensitive to the variations in A3 structure and complex composition.

### A3-Nter Fully Reconstitutes A3-WT LPL Inhibitory Effect *Ex Vivo*

To directly assess how A3 cleavage influences A3-mediated LPL inhibition, we measured total lipase activity [LPL + hepatic lipase (HL)] in pooled post-heparin plasma from the 4 genotypes. As expected, heparin increased lipase activity in all strains, with the largest increase in *Ldlr^-/-^*;*A3^-/-^* mice (Figure 3F). Post-heparin plasma was applied to a heparin affinity column to separate HL and LPL (Figure 3F, right).^41^ HL activity did not differ among strains, consistent with prior evidence that A3 does not affect HL activity.^2^ LPL activity was highest in *Ldlr*^-/-^;*A3^-/-^* mice, intermediate in *Ldlr^-/^*^-^*;A3^FL/FL^* mice and lowest in *Ldlr^-/^*^-^ mice, whereas LPL activity in *Ldlr^-/-^*;*A3^Nter/Nter^*mice was indistinguishable from *Ldlr^-/^*^-^ controls (Figure 3F, right). These results indicated that under these conditions, A3-Nter retains full LPL inhibitory capacity, while A3-FL confers only partial inhibition *in vivo*.

### A3 Cleavage Has Modest Effects on A3-Mediated Inhibition of EL

To assess the impact of A3 cleavage on EL inhibition, we measured the phospholipase A1 (PLA1) activity of post-heparin plasma. The PLA1 activity in mice expressing A3-FL and A3-Nter was comparable to that in A3-WT mice (Figure 3G), and significantly lower than in *Ldlr^-/-^*;*A3^-/-^* mice, which had a 1.5-fold increase in PLA1 activity (0.9 vs 0.6 U/mL). Thus, both A3-FL and A3-Nter retain the structural elements necessary for EL inhibition.

### Both A3 and A8 Are Present in Plasma Fractions with Maximal EL Inhibitory Activity

To further characterize the A3 complexes mediating EL inhibition, we fractionated plasma from the 4 strains on a Superdex 200 column and estimated the apparent molecular masses using calibration standards (Figure S4A). After removing the void volume, alternate fractions were immunoblotted for A3 and A8 (Figure 4B). In mice expressing A3-WT, A3-FL eluted in fractions 4-11 (158-669 kDa), A3-Nter in fractions 6-10 (158-440 kDa) and A3-Cter in nonoverlapping fractions (fractions 12-14, 44-75 kDa). A3-FL, A3-Nter, A8 and ApoA5 eluted in similar fractions across genotypes (Figure 4B and S4B). Notable, A8 eluted in the same fractions irrespective of whether the mice expressed A3, indicating that A3 is not required for its association with other plasma components.

**Figure 4.**
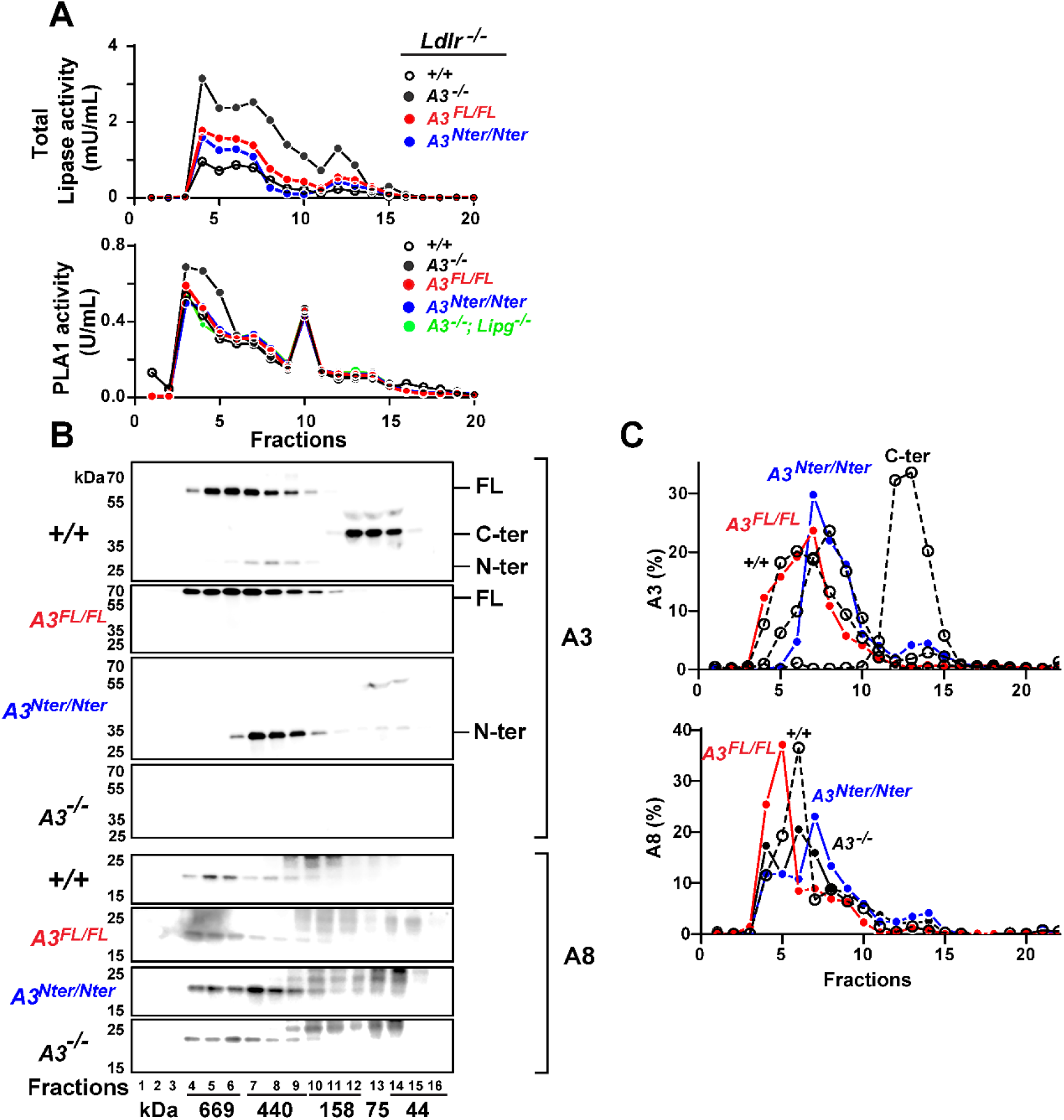
A3 and A8 co-fractionate with the highest endothelial lipase inhibitory activity. **A**, Total TG lipase and PLA1 activity in plasma fractions after size-exclusion chromatography on a Superdex 200 Increase 10/300 column. Plasma collected 15 min after heparin injection (male, n=6, 4, 6, 5 and 5/group, 8-12 wks, refed 4 h), was pooled and fractionated; after discarding the void volume (∼6 mL), fractions (330 µL/each) were collected and assayed enzymatically. **B**, Immunoblot of A3 and A8 in every other fraction (10 µL per lane). **C**, Distribution of A3-FL, A3-Nter and A3-Cter, and A8 across fractions. Band intensities were quantified by LI-COR, and the percentage of total A3 or A8 in each fraction was calculated as signal in that fraction divided by the sum of signals across all fractions. All experiments were repeated 3 times with similar results.

Since HL activity did not differ among strains (Figure 3F), we measured total lipase activity in each fraction as a proxy for intravascular LPL activity. In *Ldlr^-/-^*;*A3^-/-^* mice, lipase activity was elevated across all fractions relative to A3-expressing strains (Figure 4A, top). Both the *Ldlr^-/-^*;*A3^FL/FL^* mice and *Ldlr^-/-^*;*A3^Nter/Nter^* mice had lipase activities that were intermediate between *Ldlr^-/-^*;*A3^-/-^* mice and *Ldlr^-/-^* mice. These findings are consistent with our data showing that both A3-FL and A3-Nter partially inhibit LPL activity (Figure 3A and 3B).

The largest increase in PLA1 activity in *Ldlr^-/-^*;*A3^-/-^* mice occurred in fractions 3-5 (Figure 4A, bottom) and co-inactivation of the gene encoding EL (*Lipg*) normalized the PLA1 activity, which is consistent with prior studies,^8–10^ and confirms that the elevated PLA1 activity reflects EL activity. The PLA1 activities in *Ldlr^-/-^*;*A3^FL/FL^*and *Ldlr^-/-^*;*A3^Nter/Nter^* mice were indistinguishable from A3-WT animals, demonstrating that both A3-FL and A3-Nter fully inhibit EL at the levels of expression that occur in these animals.

The fractions exhibiting the greatest strain-dependent differences in EL activity (fractions 3-5) contained both A8 and A3 (Figure 4B), suggesting that A8 may modulate EL inhibition *in vivo*. To directly address this possibility, we inactivated *A8* in *Ldlr^-/-^* mice.

### *Ldlr^-/-^*;*A8^-/-^* Mice Have Increased Plasma Levels of A3-FL

*Ldlr^-/-^*;*A8^-/-^* mice were generated using CRISPR-Cas9 (Figure 5A), yielding a nontargeted 199 bp deletion in exon 1 of *A8,* introducing a frameshift mutation and creating a premature stop codon at position 131. Heterozygous intercrosses produced offspring at the expected Mendelian ratios (Table S4) and homozygous mutants were viable and fertile. Age-matched male *Ldlr^-/-^*;*A8^-/-^* mice had lower body (24 g vs 26.6 g, *p*=0.05) and liver (1.33 g vs 1.18 g, *p*=0.03) weights but similar liver-to-body weight ratios than *Ldlr^-/-^* controls (Figure S5A), similar to *A8^-/-^* mice.^27^

**Figure 5.**
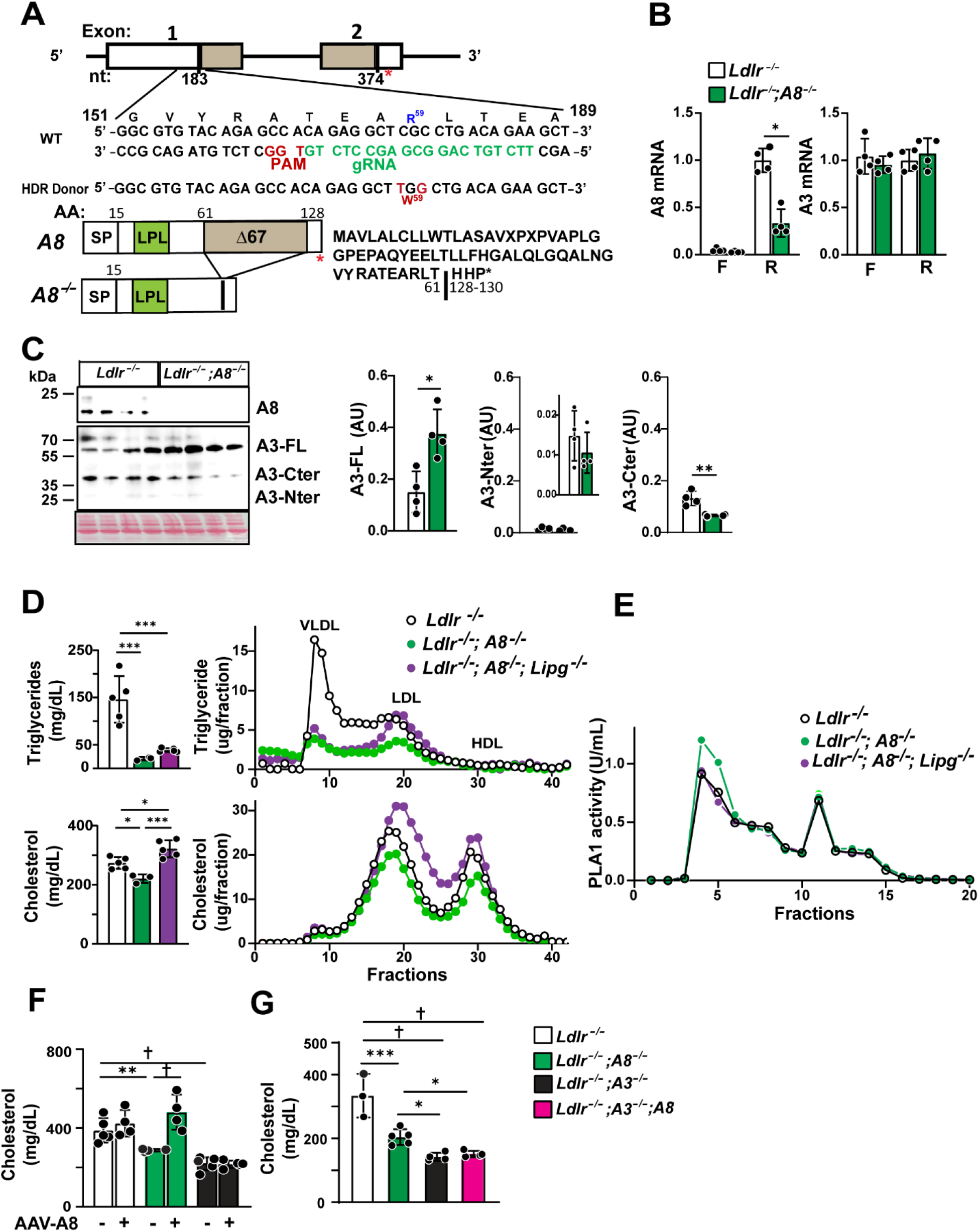
A8 deletion in *Ldlr^-/-^*mice reduced LDL-C levels and increased EL activity. **A**, CRISPR-Cas9-mediated A8 deletion in *Ldlr^-/-^* mice. A homology-directed repair template encoding the R59W substitution was introduced using a guide RNA (gRNA, green) and PAM site (red) targeting exon 1 of the *A8* gene. This editing event resulted in a 199 bp deletion (nt 184-383) and a frameshift, removing amino acids 62 to 127 and introduced a premature stop codon at residue 131, thereby abolishing A8 protein expression. **B,** Hepatic A8 and A3 mRNA in *Ldlr^-/-^*and *Ldlr^-/-^;A8*^-/-^ mice after 16 h fasting (F) and 4 h refeeding (R) (male, n=4/group, 11-13 wks). Values are normalized to the *Ldlr^-/-^* (R) group. \**p*=0.028. **C**, plasma protein levels of A8 and A3 levels in *Ldlr^-/-^*and *Ldlr^-/-^;A8*^-/-^ mice (male, n=4/group, 11-13 wks, refed 4 h). A3 was quantified by LICOR. \**p*=0.011. \*\**p*=0.003. **D,** Plasma triglyceride, cholesterol, and FPLC lipoprotein profile in *Ldlr^-/-^*, *Ldlr^-/-^;A8*^-/-^ and *Ldlr^-/-^;A8*^-/-^;*Lipg*^-/-^ mice (male, n=4, 3, and 5/group, 6-8 wks, refed 4 h). **E**, EL (PLA1) activity in *Ldlr^-/-^* and *Ldlr^-/-^;A8^-/-^* mice (male, n=4, 4, and 5/group, 11-13 wks, refed 4 h), measured as described in Fig 4A. **F**, Effect of hepatic A8 overexpression on plasma cholesterol levels in *Ldlr*^-/-^, *Ldlr*^-/-^*;A8*^-/-^, and *Ldlr*^-/-^*;A3*^-/-^ mice. 4 days after injection following a 16 h fast (male, n=5, 4, 4, 4, 5, and 5/group, 8-12 wks). **G**, Plasma cholesterol levels in *Ldlr*^-/-^, *Ldlr*^-/-^*;A8*^-/-^, *Ldlr*^-/-^*;A3*^-/-^, and *Ldlr*^-/-^*;A3*^-/-^;*A8*^-/-^ mice (male, n=3, 5, 4, and 4/group, 8-11 wks, refed 4h). Mean values were compared using Mann-Whitney test (B), parametric unpaired two-tailed student-*t* tests (C), one-way ANOVA with Tukey’s test (D), or one way ANOVA-Fisher’s LSD test (F-G) in GraphPad Prism. Data are presented as mean ± SD. \**p*<0.05, \*\**p*<0.01, \*\*\**p*<0.001, †*p*<0.0001. Statistical values were not added to the graphs when no significant differences were found. When differences were statistically significant, the exact *p* values are provided in Supplementary Table 6. All experiments were repeated at least twice with similar results.

Hepatic A8 mRNA levels, measured using primers downstream of the deletion (Table S1), were reduced by ∼70% in *Ldlr^-/-^*;*A8^-/-^* mice compared to *Ldlr^-/-^* mice (*p*=0.02) (Figure 5B, left), whereas hepatic *A3* mRNA levels did not differ between the strains (Figure 5B, right). Plasma A8 was undetectable in the *Ldlr^-/-^*;*A8^-/-^* mice (Figure 5C), while mean plasma A3-FL were 2.2-fold higher (*p*=0.01) than in *Ldlr^-/-^* mice, despite similar A3 mRNA abundance.^27^ In contrast, A3-Nter was either undetectable or present in trace amounts in both strains (Figure 5C, middle). Quantitative immunoblotting showed ∼50% lower A3-Nter and A3-Cter in the *Ldlr^-/-^*;*A8^-/-^* mice (*p*=0.003), suggesting that A8 deficiency may reduce A3 cleavage, as suggested previously,^22^ although we cannot rule out that altered A3 secretion, oligomerization, or fragment clearance may contribute to these differences.

### *Ldlr^-/-^*;*A8^-/-^* Mice Have Reduced Plasma LDL-C and Increased EL Activity

As expected, plasma TG levels were markedly lower in *Ldlr^-/-^;A8^-/-^*mice than in *Ldlr^-/-^* mice (Figure 5D, top).^27,28^ Inactivation of *Lipg*, the gene encoding EL, on this background was associated with a non-significantly higher mean levels of TG (38.7 ± 4.1 mg/dL vs 20.6 ± 3.4 mg/dL, *p*=0.71), due to an increase in the TG content of the LDL fractions (Figure 5D, top right). This may be because EL hydrolyzes VLDL-phospholipids, which generates TG-depleted LDL.^42^

Inactivation of A8 in *Ldlr^-/-^* mice was associated with a 20% reduction in mean plasma cholesterol level relative to *Ldlr^-/-^* controls (220.8 ± 14.5 mg/dL vs 274.0 ± 19.8 mg/dL, *p*=0.02; Figure 5D, bottom) due to cholesterol depletion of LDL and HDL (Figure 5D, bottom right). Inactivation of EL increased plasma cholesterol levels (Fig 5D, bottom). The mean plasma cholesterol levels in *Ldlr^-/^;A8^-/-^;Lipg^-/-^*mice exceeded that of *Ldlr^-/-^* mice (322.00 ± 29.20 mg/dL vs 273.96 ± 19.82 mg/dL, *p*=0.02), presumably due to increased remodeling and lipolysis of VLDL.^42^ These differences in plasma lipid levels occurred without alterations in hepatic TG, cholesterol or PC levels (Figure S5B).

To further evaluate the relationship between A8 deficiency and EL activity, we measured PLA1 activity in plasma fractions after size fractionation (Figure 5E). PLA1 activity peaked in fractions 4-5 in *Ldlr^-/-^;A8^-/-^* mice (Figure 5E), similar, though not identical, to the pattern seen in *Ldlr^-/-^;A3^-/-^* mice (Figure 4A) and was normalized with inactivation of *Lipg* (Figure 5E), establishing that EL expression is required for the LDL-C-lowering effect of A8 deficiency.

### A8 Requires A3 to Elevate Plasma Cholesterol Levels

We overexpressed A8 in *Ldlr^-/-^;A8^-/-^* and *Ldlr^-/-^;A3^-/-^* mice. A8 reconstitution normalized plasma cholesterol levels in *Ldlr^-/-^;A8^-/-^* mice, but not in *Ldlr^-/-^;A3^-/-^* mice (Figure 5F). Similar results were seen for plasma TG levels (Figure S6A and S6B). Thus, A3 expression is required for the lipid-elevating effects of A8 expression. Inactivating A8 in *Ldlr^-/-^;A3^-/-^* mice produced no further reduction in plasma cholesterol or TG levels (Figure 5G and S6C). The effect of inactivating A8 on plasma cholesterol level was similar, irrespective of whether the mice expressed A3-WT, A3-FL or A3-Nter (Figure S6D).

Collectively, these data show that the cholesterol-elevating activity of A8 is dependent on the presence of A3, establishing A3 as an essential cofactor for A8-medicated regulation of plasma cholesterol levels.

### A8 Enhances A3’s Inhibition of EL

To clarify how A8 increases A3-mediated EL inhibition, we compared the effect of A3 to A3 plus A8 on EL activity *in vitro*. Recombinant V5 tagged A3 and A3 plus A8 were expressed in QBI 293A cells, enriched by V5 immunoprecipitation, quantified by ELISA, and verified by immunoblotting (Figure 6A). Conditioned medium expressing EL was incubated with A3 or A3/A8 and EL activity was measured using a commercial PLA1 assay.^30^ At 12.5 ng of A3, the A3 plus A8 complex inhibited EL ∼ 20% more than A3 alone (Figure 6B, *p*=0.003), with the difference diminishing at higher A3 concentrations. These results indicate that A3/A8 complex is a more potent EL inhibitor than A3 alone under these conditions.

**Figure 6.**
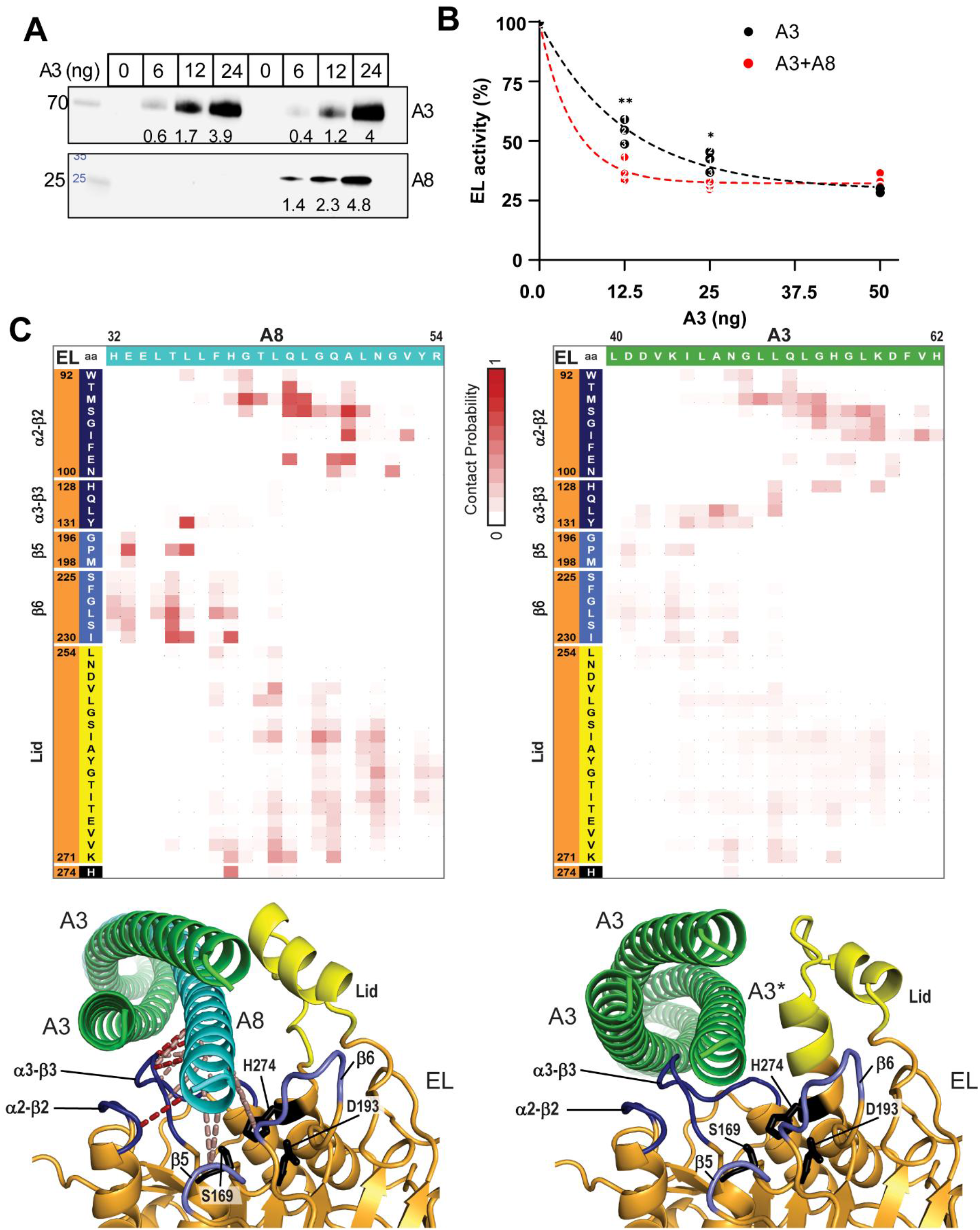
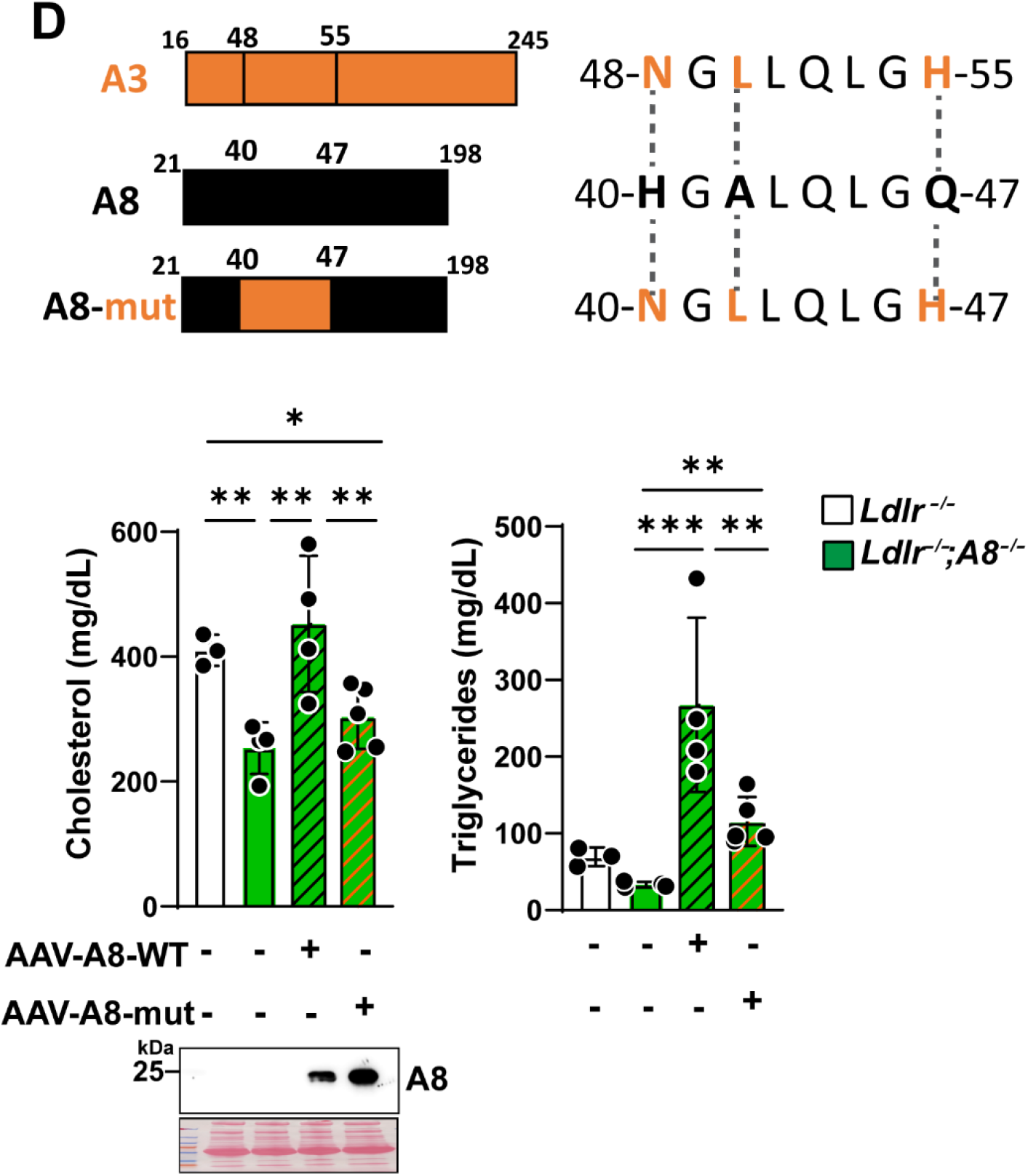
A8 modulates EL activity through A3. **A**, A3 concentrations were determined by human A3 ELISA and confirmed by immunoblotting; A3 and A8 band intensities were quantified using LI-COR. **B**, EL phospholipase activity after incubation with the indicated amounts of A3 or A3/A8 as described in Supplementary methods. The data fitted into a mono-phased decay curve using Graphpad Prism. \*\**p*=0.003. \**p*=0.03 **C**, AlphaFold structure models of EL in complex with A3/A8. Top: AlphaFold contact probability matrices between residue pairs are colored in heatmap as labeled. Color scale bar: 0 to 1 with units of contact probability. Sequence ranges for EL (orange, rows) are labeled according to structure features and depicted with sequences from either A8 (cyan, columns, left panel) or corresponding A3 (green, columns, right panel). Bottom: EL (orange cartoon) is colored by structure features depicted in the heatmap: active site (black), key interacting loops (blue shades) and lid (yellow) and interacts with A3_2_ (green cartoon) complexed with A8 (cyan cartoon, left) or A3* (green cartoon, right). Residue pairs with high contact probability are connected by colored dash lines: (> 0.7, red or between 0.6 and 0.7, salmon). **D**, Top: A8-mut was generated by introducing A3-like residues into mouse A8-WT (H40N, A42L, and Q47H). Bottom: Effect of hepatic A8 mutant overexpression on plasma cholesterol levels in *Ldlr*^-/-^*;A8*^-/-^ mice. 6 days after injection following a 16 h fast (male, n=3, 4, 4 and 5/group, 5-9 wks). A8-WT and mutant expression were determined by Immunoblot. Mean values were compared using paired two-tailed *t* tests (B) or one way ANOVA-Fisher’s LSD test (D) in GraphPad Prism. Data are presented as mean ± SD. \**p*<0.05, \*\**p*<0.01, \*\*\**p*<0.001. Exact *p* values in Fig 6D were provided in Supplementary Table 6. All experiments were repeated at least twice with similar results.

We used AlphaFold to generate model complexes between EL and a homotrimer of A3 (A3_3_) or a heterotrimer containing two A3 plus one A8 subunit (A3_2_A8). Although global interaction confidence for full-length complexes with models is marginal (Table S5), the EL:A8 interface in the A3_2_A8 model contained high confidence co-evolution-derived contact probabilities, which identified several interaction hotspots within the N-terminal portion of A8 (Figure 6C, upper left). In the EL:A3_3_ model, the corresponding residues showed weaker contact probabilities (Figure 6C, upper right).

The predicted EL structure closely resembled published models of LPL (Figure S7), including the lid and surrounding loops.^43^ In the EL:A3_2_A8 model (Figure 6C, lower left), the N-terminal helix of A8 lies adjacent to the EL active site, and engages with the catalytic histidine (black). The A8 residues with the highest probability interact with residues from EL β2-α2 and β3-α3 loops (Figure 6C, lower left, dark blue), which are analogous to LPL loops that were protected from deuterium exchange upon incubation with A3/A8.^43^ Additional loops following the EL β5 and β6 strands (Figure 6C, lower left, light blue) include residues with intermediate confidence scores that promotes the positioning of the A8 helix near to the active site.

In the EL:A3_3_ model (Figure 6C, lower right), the primary A3 helix occupies a similar overall binding orientation as A8 in the EL:A3_2_A8 complex, but is displaced laterally away from the active site, consistent with lower interface confidence (Figure 6C, upper right). Collectively, the coevolution-derived inter-residue contact patterns at the EL interaction surface support a stronger and more focused interaction between EL and A8 than between EL and A3, providing a structural rationale for the enhanced EL inhibition by the A3_2_A8 complex.

To test this hypothesis, we introduced A3-like residues into mouse A8 (H40N, A42L, and Q47H) and injected mutagenized AAV-A8 (AAV-A8-mut) into A8-deficient mice (Figure 6D). Compared to AAV-A8-WT, the mutant A8 failed to increase cholesterol levels despite higher protein levels. In addition, the plasma TG-elevating effect was attenuated in mice expressing the mutant A8 compared with WT A8, suggesting that this A8 motif contributes to its inhibition of LPL and EL

## Discussion

In this study, we show that A3 cleavage and A8 act together *in vivo* to modulate EL as well as LPL, thereby maintaining cholesterol and TG homeostasis in *Ldlr^-/-^* mice. A major new finding is that inactivation of A8 in *Ldlr^-/-^* mice lowers cholesterol by ∼20%. This hypocholesterolemic effect requires expression of both A3 and EL, indicating that A8 contributes to A3-mediated inhibition of EL *in vivo*, in addition to its established role in inhibiting LPL. In our study, the highest level of LPL inhibition was observed when A3-FL and A3-Nter were present together with A8, whereas the highest EL inhibition depended on A8 but not on A3 cleavage, revealing partially overlapping yet nonredundant functions of A3 isoforms and A8 in intravascular lipoprotein metabolism.

The physiological importance of A3 is underscored by its strong conservation across vertebrates, whereas the furin cleavage site in A3 is conserved only in mammals.^20^ We provide *in vivo* evidence that optimal A3-mediated LPL inhibition occurs when both A3-FL and A3-Nter are expressed (Figure 3B). The juxtaposition of strong A3 conservation with mammal-restricted furin cleavage supports a model in which regulated A3 proteolysis has evolved to tune the balance between a relatively stable full-length pool and a shorter lived, N-terminal pool of A3 to regulate clearance of circulating TG. Our *in vivo* data further suggest that these two A3 isoforms, together with A8, form a trimeric A3-FL/A3-Nter/A8 complex to inhibit LPL. Consistent with this model, selective blockade of A3 cleavage disproportionately impaired LPL inhibition compared with EL inhibition.

Neither A3 cleavage, nor the presence of A3-Cter appears to be essential for survival, at least under the conditions studied. The FBG domain, which is present in ANGPTs and other ANGPTL family members, comprises A, B, and P subdomain arranged around a central β sheet.^44^ The P subdomain contains several surface loops that in ANGPTs bind vascular receptors. The P subdomain of A3-Cter can interact with the integrin receptor avβ3 and support endothelial adhesion and signaling in cultured cells, but the physiological significance of the interaction remains unproven.^45^ The FBG domain of A4 stimulates adipocyte lipolysis,^46^ yet no analogous activity has been demonstrated for A3-Cter. Other than effects on plasma lipid levels, we observed no differences between mice expressing no A3-Cter (*Ldlr^-/-^*;*A3^Nter/Nter^*and *Ldlr^-/-^*;*A3^-/-^* mice) and lacking any free A3-Cter (*Ldlr^-/-^*;*A3^FL/FL^* mice). Thus, despite strong evolutionary conservation of the FBG domain in A3, we found no adverse consequences of its absence in these animal models. We cannot rule out the possibility that phenotypes might emerge under metabolic stress, tissue injury, or other environmental challenges.

A key finding of this work is that A8 inactivation lowers plasma LDL-C and HDL-C in *Ldlr^-/-^* mice, and that this hypocholesterolemic effect requires both A3 and EL. This result indicates that A8 contributes to A3-mediated inhibition of EL *in vivo*. Only modest differences in plasma cholesterol levels were apparent in *Ldlr^-/-^*;*A3^FL/FL^*, *Ldlr^-/-^*;*A3^Nter/Nter^* and *Ldlr^-/-^* mice, suggesting that A3-Cter is not required to stabilize the interaction between A3 and EL in the presence or absence of A8 *in vivo*.

Our *in vivo* genetic data indicates that A8 is required for full A3-EL inhibition, at least in the absence of LDLR. These results differ from prior studies using conditioned media where A8 did not alter the binding or the inhibition of EL by A3,^30^ but support work using purified-components.^47^ Our results are consistent with population-based studies in which higher plasma ABCL levels and coronary heart disease (CHD) risk correlate better with levels of circulating A3/8 than A3 alone,^48^ revealing a physiological requirement for A8 in regulating LDL-C levels.

Our AlphaFold-based interface predictions identified localized, binding regions that are predicted to mediate A8-EL interactions. The structural model of the A3_2_A8 trimer bound to EL resembles the interaction surface defined by hydrogen-deuterium exchange experiments with LPL^43^ and suggests that A8 provides a higher-confidence, higher-affinity interface with EL than A3_3_. AlphaFold contact probabilities between the A8-Nter helix and EL exceed those for A3, supporting a division of labor in which A3 functions primarily as a stabilizing scaffold within the heterotrimer, whereas A8 furnishes key contact residues that optimize EL engagement. Within this framework, EL inhibition is achieved most efficiently by the A3/A8 complex irrespective of A3 isoform, whereas LPL inhibition is more sensitive to the balance between A3-FL and A3-Nter *in vivo*.

Despite A3 deficiency exerting substantial effects on plasma LDL-C, our perfusion studies in mice showed that it did not alter the rate of ABCL secretion or the lipid content of the newly secreted VLDL particles. This differs from human kinetic data, which reported a reduction in liver-derived TRL-apoB and TG production and an increase in TRL clearance.^49^ In that study, the authors did not sample the newly secreted VLDL directly from the liver, as we have done. Rather, they have used circulating VLDL, that by definition is the least efficiently cleared. Their methodology does not detect the particles that are secreted and then rapidly removed by the liver before entering the peripheral circulation. Since these data were obtained using fundamentally different experimental approaches, we cannot determine from the available data whether the discrepancy in results reflects species differences in hepatic TRL production, though we think it more likely reflects methodological differences. These data, together with prior work showing that A3 inactivation lowers LDL-C independently of LDLR, LRP1, SCARB1, syndecan-1, and ApoE in mice,^9,21,36^ support the existence of a yet-to-be-defined hepatic uptake pathway that preferentially clears newly-secreted ABCLs from the circulation. The reason for the marked reduction in rate of TG appearance in plasma^36^ in A3-deficient animals after LPL inhibition remains unclear, but these data are compatible with a scenario in which nascent ABCLs are rapidly removed by the liver before entering the peripheral circulation.^50^

A major challenge for A3-targeted lipid-lowering is the near-complete neutralization of A3 appears necessary to achieve clinically meaningful LDL-C lowering in humans. Achieving very low A3 activity levels is challenging and currently requires intravenous administration of anti-A3 antibodies due to high circulating levels of A3. Alternative anti-A3 strategies include antisense oligonucleotide (e.g., vupanorsen) and siRNA-based approaches (e.g., zodasiran, solbinsiran).^15,16,17^ Development of vupanorsen was halted due to increased hepatic TG and liver enzymes, whereas siRNA-based agents, though subcutaneously administered and generally being well-tolerated, have more modest LDL-lowering effects than those observed in A3-deficient individuals. CRISPR–Cas9 targeting A3 offers a “one-and-done” strategy, but concerns persist regarding long-term safety and efficacy.^51^ An alternative and potentially more effective approach is to simultaneously target A3 and its cofactor A8. In a recent phase 1 trial using a mAb that disrupts the interaction between A3/A8 and LPL resulted in a 32% reduction in LDL-C as well as a 70% reduction in TG in patients with mixed hyperlipidemia.^52^ It is plausible that a combination of A3 and A8 inhibition, or agents engineered to disrupt A3/A8-EL interaction will prove highly effective in lowering plasma LDL-C levels than A3-directed monotherapy.

Together, these findings support dual A3-A8 inhibition as a strategy to achieve robust cholesterol as well as TG lowering and highlight the A3/A8-LPL/EL axis as a rational focus for next-generation therapeutics.

## Acknowledgements

We thank Lisa Beatty, Linda Donnelly, Tommy Hyatt, Fang Xu, and Christina Zhao, for their excellent technical support. We thank Drs. Qinli Hu and Haili Cheng for helpful discussions. We thank Andrew Lemoff and Xuemei Luo in the UTSW Proteomics Core for protein quantification.

## Disclosures

### AI Use Disclosures

Artificial intelligence-assisted tools were used for structural modeling and content condensing to meet word limits.

## Nonstandard Abbreviations and Acronyms

AAV: adeno-associated virus
A3 ANGPTL3: angiopoietin-like protein 3
A4 ANGPTL4: angiopoietin-like protein 4
A8 ANGPTL8: angiopoietin-like protein 8
ASO: antisense oligonucleotide
CHD: coronary heart disease
Cter: C-terminal fragment
FL: full-length protein
Nter: N-terminal fragment
WT: wildtype
ABCLs: apolipoprotein (Apo)B containing lipoproteins
ANGPTs: angiopoietins
ANGPTLs: angiopoietin-like proteins
ASCVD: atherosclerotic cardiovascular disease
EL: endothelial lipase
ELISA: enzyme linked immunoabsorbent assay
FBG: fibrinogen-like domain
GPIHBP1: glycosylphosphatidylinositol-anchored high-density lipoprotein binding protein 1
HDL: high density lipoprotein
HSD: high-sucrose diet
HL: hepatic lipase
LDL: low density lipoprotein
LDL-C: LDL-cholesterol
LDLR: LDL receptor
LPL: lipoprotein lipase
LRP1: LDL receptor related protein 1
PC: phosphatidylcholine
PLA1: phospholipase A1
siRNA: small interfering RNA
SRM/MS: Selected reaction monitoring mass spectrometry
TG: triglyceride

