## Supplementary material for "Dual Control of LDL-cholesterol Levels by ANGPTL3 and ANGPTL8"

**Short title**: ANGPTL3 and ANGPTL8 in Cholesterol Metabolism

University of Texas Southwestern Medical Center (UTSW)

Dallas, TX 75390-9046

**Expanded Materials and Methods**:

- Supplementary methods
- Supplementary table 1-6
- Supplementary figure 1-7

### Supplementary Methods

**Table1. Materials: Reagents used in the experiments described in this paper.**

| REAGENT | SOURCE | IDENTIFIER |
|---|---|---|
| **Antibodies (Ab)** | | |
| Donkey Anti-goat IgG $2^{o}$ Antibody (Ab) | Jackson ImmunoResearch | 705-035-003 |
| Donkey Anti-mouse IgG $2^{o}$ Ab | Jackson ImmunoResearch | 715-035-150 |
| Goat Anti-mouse IgG, Fcy fragment-specific | Jackson ImmunoResearch | 115-035-071 |
| Goat Anti-rabbit IgG $2^{o}$ Ab | Jackson ImmunoResearch | 111-035-144 |
| Goat Anti-ANGPTL3 (A3) polyclonal (p)Ab | Bio-Techne | AF136 |
| Mouse Anti-ANGPTL8 (A8) monoclonal (m)Ab | In-house, as described[53] | 19H4 |
| Mouse Anti-V5 mAb | Invitrogen | 46-1157 |
| Rabbit Anti-A3 mAb | In-house (see below) | 2G5 |
| Rabbit Anti-ApoA5 pAb | US Biological | 139307 |
| Rabbit Anti-ApoB pAb | Abcam | Ab20737 |
| Rabbit Anti-ApoE mAb | Abcam | Ab183596 |
| Rabbit Anti-Calnexin pAb | Enzo | ADI-SPA-860 |
| Rabbit Anti-Flag pAb | Millipore Sigma | F7425 |
| Rabbit Anti-GFP mAb | Cell Signaling | 2956S |
| **Bacterial Strains** | | |
| *E. coli* BL21 (DE3) Competent Cells | Thermo Fisher Scientific | EC0114 |
| **Buffers** | | |
| Dulbecco′s Phosphate Buffered Saline (DPBS) | Millipore Sigma | D8537 |
| Laemmli SDS Sample Buffer, Reducing 6x | Thermo Fisher Scientific | J61337.AC |
| Laemmli SDS Sample Buffer, Non-reducing 6x | Thermo Fisher Scientific | J60660.AC |

| | | |
|---|---|---|
| NativePAGE 20x Cathode Buffer Addictive | Invitrogen | BN2002 |
| NativePAGE™ Sample Prep Kit | Invitrogen | BN2008 |
| NativePAGE 20x Running Buffer | Invitrogen | BN2001 |
| Phosphate Buffered Saline (PBST) | Millipore Sigma | P3563 |
| Phosphate Buffered Saline (10x PBS) | Millipore Sigma | 6505-4L |
| Sodium Chloride Irrigation (0.9%), USP | Baxter | 2F7123 |
| Trans-Blot Turbo 5x Transfer Buffer | Bio-Rad | 10026938 |
| UltraPure™ 0.5M EDTA, pH 8.0 | Thermo Fisher Scientific | 15575038 |
| **Cell Lines** | | |
| QBI-HEK293A | Obiogene, formerly Quantum Biotechnologies | QBI-293A-2 |
| **Chemicals** | | |
| Acetic Acid | JT Baker | 9515-03 |
| Bovine Serum Albumin (BSA) | Sigma | A3803 |
| FuGENE® 6 Transfection Reagent | Promega | E2692 |
| Glycerol | Millipore Sigma | G9012 |
| Heparin Sodium Salt | Sigma | H3393-500KU |
| Imidazole | Sigma-Aldrich | 56750 |
| Isopropyl β- d-1-Thiogalactopyranoside (IPTG) | Millipore Sigma | I6758 |
| Kanamycin Sulfate | Millipore Sigma | K1377 |
| NativeMark™ Unstained Protein Standard | Thermo Fisher | LC0725 |
| Ni-NTA Agarose | Qiagen | 30230 |
| Potassium Bromide (KBr) | Sigma | P0838 |
| Protease Inhibitor Cocktail (cOmplete™ Mini EDTA-free) | Millipore Sigma | 11836170001 |

| | | |
|---|---|---|
| Sodium Hydrochloride (NaCl) | Sigma-Aldrich | S3014 |
| Tris Hydrochloride (Tris HCl) | Sigma-Aldrich | 10812846001 |
| cis-4,7,10,13,16,19-Docosahexaenoic Acid | Sigma | D2534-1G |
| Linoleic Acid | Sigma | L1376-10G |
| γ-Linolenic Acid | Sigma | L2378-1G |
| Oleic Acid | Sigma | O1008-5G |
| Palmitoleic Acid | Sigma | P9417-1G |
| Palmitic Acid | Sigma | P5585-10G |
| Stearic Acid | Sigma | S4715-1G |
| **Kits** | | |
| Cholesterol Quantitation Kit | Sigma Aldrich | MAK043 |
| EnzChek™ Phospholipase A1 (PLA1) Assay Kit | Invitrogen | E10219 |
| Gel Filtration HMW Calibration Kit | Cytiva | 28403842 |
| High Sensitivity Triglyceride (TG) Assay Kit | Sigma Aldrich | MAK264 |
| Human A3 ELISA kit | Bio-Techne | DANL30 |
| Lipoprotein Lipase (LPL) Activity Assay | Cell Biolabs | STA-610 |
| Mouse ApoB-100 ELISA Kit | Abcam | ab230932 |
| Mouse A3 ELISA Kit | Antibodies online | ABIN1884517 |
| Phospholipid Quantification Assay Kit | Millipore Sigma | CS0001 |
| Phospholipids C Reagent | Wako life Sciences | 997-01801 |
| Pierce BCA Protein Assay Kit | Thermo Fisher Scientific | 23224 |
| Power SYBR Green PCR Master Mix | Applied Biosystems | 4368708 |
| Total Cholesterol Reagents | Thermo Fisher | TR13421 |
| Triglycerides Reagent | Thermo Fisher | TR22421 |
| TaqMan Reverse Transcription Reagents | Applied Biosystems | N8080234 |

| | | |
|---|---|---|
| RNeasy Plus Universal Mini Kit | Qiagen | 73404 |
| V5-tagged Protein Purification Kit Ver.2 | Cosmo Bio | MBL-3317 |
| **Medium** | | |
| Dulbecco′s Modified Eagle Medium (DMEM) with L-Glutamine, 4.5g/L Glucose | Corning | 10-0130-CV |
| Fetal Bovine Serum (FBS) | Corning | 35-015-CV |
| Luria Broth (LB) | Fisher Scientific | BP1426-500 |
| Opti-MEM™ Reduced Serum Medium | Gibco | 11058-021 |
| **Oligonucleotides** | | |
| See SI Appendix, Table S1 | | |
| **Recombinant DNA** | | |
| pcDNA3.1 | Lab | |
| pET28a | Novagen | 69864 |
| **Recombinant Proteins** | | |
| Human-A3-V5-6xHis | Lab | |
| Human-A8-V5-6xHis | Lab | |
| Human-EL-Myc-Flag | Lab | |
| Mouse-A3 (aa17-224)-6xHis | Lab | |
| **Software and Algorithms** | | |
| Image Studio Lite v5.2 | LI-COR | Version 5.2.5 |
| GraphPad Prism 10 | GraphPad | Version 10.0.0 |

| | | |
|---|---|---|
| SnapGene | SnapGene | Version 6.0.2 |
| **Virus** | | |
| AAV8-TBG-GFP | Vector Biolabs | QB63528 |
| AAV8-TBG-mA8-Flag | Vector Biolabs | QB63094 |
| AAV8-TBG-mA8 (mutant)-Flag | Vector Biolabs | Q-0163-1 |
| **Other** | | |
| Amicon Ultra Centrifugal Filter (10 kDa MWCO) | Millipore Sigma | UFC8010 |
| AKTA Pure Chromatography System | Cytiva | 29018224 |
| Chow Diet (rodent) 16% Protein | Teklad Global | 2016 |
| 4–15% Criterion™ TGX™ Precast Midi Protein 26-well Gel | Bio-Rad | 5671085 |
| 4–15% Criterion™ TGX™ Precast Midi Protein 18-well Gel | Bio-Rad | 5671084 |
| 4–15% Criterion™ TGX™ Precast Midi Protein Gel (Prep+2 well) | Bio-Rad | 5671082 |
| HisTrap HP, 5 ml | Cytiva | 17524802 |
| Immobilon®-PSQ PVDF 0.2 μm Membrane | Millipore Sigma | ISEQ00010 |
| Instant Nonfat Dry Milk | Nestle | |
| Millex-HV Syringe Filter Unit (0.45 μm, PVDF membrane) | Millipore Sigma | SLHV033RS |
| MP Biomedicals™ Fat Free Diet, Pellet (high sucrose diet) | Thermo Fisher Scientific | ICN90168310 |
| NanoDrop™ 8000 Spectrophotometer | Thermo Fisher Scientific | ND-8000-GL |
| NativePAGE™ Bis-Tris Mini Protein Gels (3-12%) 1.0 mm | Thermo Fisher Scientific | BN2011BX10 |
| Open-Top Ultra-Clear Tube (13.2 mL) | Backman | 344059 |
| Superose 6 Increase 10/300 GL Column | Cytiva | 29091596 |

| | | |
|---|---|---|
| Superdex 200 Increase 10/300 GL Column | Cytiva | 28990944 |
| SuperSignal™ West Pico PLUS Chemiluminescent Substrate | Thermo Fisher Scientific | 34577 |
| Trans-Blot Turbo RTA Midi 0.2 µm Nitrocellulose Transfer Kit | Bio-Rad | 1704271 |

**Mice**

Mice were housed in a controlled environment (12-h light/12-h dark cycle) at ambient temperature of 23 ± 1°C, 60-70% humidity). For fasting/refeeding experiments, mice were entrained for at least 3 days. The mice were fasted overnight (5:00 PM-9:00 AM) and fed during the day (9:00 AM-5:00 PM). Experiments were performed at 9:00 AM (fasted) or 1:00 PM (4 h after refeeding).

$A3^{-/-}$ mice were generated by replacing the gene with a LacZ-Hyg cassette using homologous recombination. The mice were backcrossed with C57BL/6J mice as described.[38] $Ldlr^{-/-};A3^{FL/FL}$ mice were generated as described in the legend to Figure 2. For $Ldlr^{-/-};A3^{Nter/Nter}$ mice, a stop codon followed by a polyadenylation (polyA) tail was inserted at residue 225. $F_0$ founders were crossed with $Ldlr^{-/-}$ mice to generate heterozygous offspring ($F_1$), which were intercrossed to generate mice homozygous for the mutation. $Ldlr^{-/-};A3^{FL/-}$ and $Ldlr^{-/-};A3^{Nter/-}$ mice were generated by crossing $Ldlr^{-/-};A3^{FL/FL}$ mice and $Ldlr^{-/-};A3^{Nter/Nter}$ with $Ldlr^{-/-};A3^{-/-}$ mice, respectively. $Ldlr^{-/-};A3^{FL/Nter}$ heterozygous mice were obtained by intercrossing $Ldlr^{-/-};A3^{FL/-}$ and $Ldlr^{-/-};A3^{Nter/-}$ mice. Triple knockout $Ldlr^{-/-};A8^{-/-};Lipg^{-/-}$ mice were produced by crossing $Ldlr^{-/-};A8^{-/-}$ mice with $Ldlr^{-/-};Lipg^{-/-}$ mice following by intercrossing the double- heterozygous offspring. Similarly, $Ldlr^{-/-};A3^{-/-};A8^{-/-}$ mice were generated by crossing $Ldlr^{-/-};A8^{-/-}$ mice with $Ldlr^{-/-};A3^{-/-}$ mice. $Ldlr^{-/-};A3^{FL/FL}$ and $Ldlr^{-/-};A3^{Nte/Nter}$ mice were crossed with $Ldlr^{-/-};A8^{-/-}$ mice. The double-heterozygous offspring were then intercrossed to generate $Ldlr^{-/-};A3^{FL/FL};A8^{-/-}$ and $Ldlr^{-/-};A3^{Nter/Nter};A8^{-/-}$ mice.

Genomic DNA was extracted from mouse tails and analyzed by PCR for genotyping. Primer sequences are listed in Table S1.

For the mouse experiments, no formal *a priori* sample sizes were calculated. Sample sizes were based on prior experience with similar mouse studies, the expected effect sizes, and the availability of the required genotypes. The number of animals used in each experiment is indicated in figure legends.

For experiments comparing genotype-defined groups, randomization was not used. For intervention experiments within the same genotype (Figure 3E, 5F and 6D), animals from the same cage/litter were randomly distributed between groups to minimize allocation bias. Experimental procedures were performed in a balanced order to minimize potential bias.

**Liver Perfusion and VLDL Isolation**

Livers from ad libitum fed mice were perfused exactly as described.[54] Mice were anesthetized with ketamine/xylazine (4U/g body weight) followed by an intraperitoneal injection of heparin (100 U). Perfusions were conducted for 60 min in a nonrecirculating mode using oxygenated buffer containing free fatty acids (FFA, 0.8 mmol/L) bound to albumin. Following each perfusion, VLDL-enriched fractions were isolated from perfusates by density gradient ultracentrifugation (d=1.006 g/mL using KBr). For heparin-treated samples, VLDL was isolated as described previously.[55] For non-heparin treated samples, perfusates (12 mL) were subjected to centrifugation (40,000 rpm in TH641 swinging bucket rotor, 18 h at 4°C) in ultra-clear open-top tubes. The top 1 mL was collected, and proteinase inhibitors were added before the samples were stored at -80°C.

Immunoblotting of VLDL proteins was performed as described.[55] VLDL-lipids (PC, cholesterol, TG) were quantified fluorometrically using kits (see Table 1).[55]

**Protein Expression and Purification for Antibody Generation**

A cDNA encoding mouse A3 (residues 17-224) was cloned into pET28(+) and the sequence was verified prior to the plasmid being introduced into *E. coli* BL21 cells. Kanamycin-resistant colonies were expanded in LB-kanamycin (1 L) to an $OD_{600}$=0.8, and then IPTG (1 mM) was added to induce expression, and the cells were grown for 3 h at 37°C.

Cells were pelleted (3325*g* for 30 min at 4°C), resuspended in 10 mL of buffer A (20 mM Tris HCl, 500 mM NaCl, 5 mM imidazole, 10% glycerol, pH 8.0) and lysed by sonication (3 sec on, 4 sec off, 5 min total) on ice. The lysates were clarified by centrifugation (15,000 rpm, 30 min, 4°C). The His-tagged proteins were purified using Ni-NTA resin (1 mL/L). Cell lysates were incubated with Ni-NTA resin equilibrated in buffer A at 4°C for 1 h on a shaker (200 rpm), washed 3x with buffer B (20 mM Tris HCl, 500 mM NaCl, 25 mM imidazole, 10% glycerol, pH 8.0), then eluted in buffer C (20 mM Tris HCl, 300 mM NaCl, 300 mM imidazole, 5% glycerol, pH 8.0). Purified A3-Nter (aa 17-224) was concentrated using a membrane filter (10-kDa) and then quantified by absorbance at A280, aliquoted (1mg/mL), snap-frozen, and stored at -80°C for antibody generation.

**Immunoassays**

Immunoblotting was performed after protein separation by SDS-PAGE (4-15%) as described previously.[52] Mouse plasma was diluted 1:5 in 0.9% NaCl and heated at 95°C for 5 min in Laemmli reducing sample buffer before electrophoresis. A total of 10 μL of diluted plasma was loaded per lane. To detect A3, plasma samples were prepared in non-reduced Laemmli buffer, and 5 μL was loaded per lane unless indicated otherwise in figure legends.

Membranes were incubated overnight at 4°C with the indicated primary antibodies (1:1000), except for the anti-A8 mAb (19H4, 0.6 mg/mL), which was used at a dilution of 1:250. The 2° antibodies (1:5000 dilution) were incubated with the membranes for 1 h at room temperature. A complete list of antibodies used for immunoblotting is provided in Table 1.

Native protein complexes were separated on NativePAGE (3-12%) Bis-Tris gels according to the manufacturer's instructions. Plasma from male mice (4-5/per group, 8-12 wks, refed for 4 h) was fractionated by Superdex 200 size-exclusion chromatography. Fractions containing immunoreactive A3 were pooled, concentrated by centrifugation using 10 kDa cut-off filters (5000*g* for 20 min at 4°C), and prepared in 1x NativePAGE sample buffer before electrophoresis. A total of 30 µL was loaded per lane, except for the sample from mice expressing A3-FL, for which 10 µL was loaded because of higher protein abundance.

Fractionated plasma was separated by native PAGE (first dimension) at room temperature for 1.5 h at 150 V using prechilled light blue cathode buffer in the upper chamber and 1× NativePAGE anode buffer in the lower chamber. Proteins were transferred to 0.2 µm PVDF membranes using 1× Turbo blot Transfer Buffer, washed with 8% acetic acid for 15 min, blocked in 5% nonfat milk for 1 h, and incubated overnight at 4°C with anti-A3 mAb (2G5). Lane strips were excised from the nondenaturing gel and subjected to SDS-PAGE under denaturing conditions in a second dimension. Proteins were then transferred to 0.22-µm nitrocellulose membranes and immunoblotted with antibodies against A3 (2G5) and A8 (19H4).

Immunoblots were imaged using an Odyssey FC Imager (LI-COR) and the band intensities were quantified with Image Studio Lite Ver 5.2 (LI-COR).

#### Quantitative Real-Time PCR

Total RNA was extracted from liver samples using an RNA extraction kit according to the manufacturer's instructions. RNA concentration and purity were determined using a NanoDrop spectrophotometer, and RNA integrity was assessed by agarose gel electrophoresis. First-strand cDNAs were synthesized from 2 μg total RNA using random hexamer primers and reverse transcription reagents (see Table 1).

Real-time PCR was performed using an Applied Biosystems real-time PCR system with SYBR Green (Table 1). Each reaction was carried out in a 20 μL volume containing 20 ng cDNA, 167 nM primers, and 10 μL SYBR Green PCR Master Mix, in duplicate.[56] Glyceraldehyde-3-phosphate dehydrogenase (*Gapdh*) mRNA and 18S ribosomal RNA served as the internal normalization control.

**Adeno-Associated Virus Infection of Mice**

Adeno-associated virus serotype 8 (AAV8) vectors expressing either GFP (control), mouse A8 or mutant mouse A8 under the control of the TBG promoter were administered by tail-vein injection at a dose of $1\times10^{11}$ genome copies in 200 μL saline per mouse using a 1-mL syringe with a 30G×½" needle. After injection, mice underwent at least 3 fasting/refeeding cycles and were killed after the final fasting period.

**Lipid Analysis and Lipoprotein Profiling**

Hepatic lipids were extracted from liver (~50 mg) using Folch solution.[57] Levels of hepatic and circulating TG, cholesterol and PC were measured enzymatically as described.[22] Hepatic lipid levels were normalized to wet tissue weight (mg/g).

Plasma lipoprotein particles were size-fractionated by fast performance liquid chromatography using a Superose 6 Increase 10/300 GL column. Cholesterol and TG content of each fraction was measured enzymatically as described.[22]

**LPL and EL Activity Assays**

Heparin (1 U/g) was injected intravenously after 4 h of refeeding. Plasma was collected 15 min post-injection, isolated by centrifugation (2000*g* for 10 min at 4°C), and snap-frozen as described previously.[38] Total lipase activity and PLA1 activity in plasma were measured using enzymatic kits (Table 1) according to the manufacturers' instructions.[30,52]

To separate hepatic lipase (HL) and lipoprotein lipase (LPL), post-heparin plasma (250 µL/group) was fractionated on a 1 mL heparin-affinity column as described.[41,52] TG lipase activity in each fraction (20 µL/fraction) was measured using a TG lipase activity kit according to manufacturer's instructions. Additionally, plasma (350 µL/group) was fractionated on a Superdex 200 size exclusion column, and odd fractions (10 µL/fraction) were analyzed for both TG lipase and PLA1 activities as described.[30,52]

**Cell Lines and Transfections**

QBI-HEK293A cells were grown in 10-cm dishes in complete medium [high sucrose DMEM plus 5% FBS] at 37 °C in 8.8% $CO_2$. Cells were transiently transfected with cDNAs encoding human A3, A8, or EL using FuGENE 6 according to the manufacturer's instructions. Conditioned medium was collected 72 h post-transfection, filtered through a 0.45 µm membrane to remove cell debris, and concentrated using a 10-kDa membrane filter (5,000*g* for 15 min at 4°C). The concentrated medium was used immediately or snap-frozen and stored at -80°C.

**Enzyme-Linked Immunosorbent Assay (ELISA)**

Plasma A3 was measured using a sandwich ELISA according to the manufacturer's instructions. In this assay, the capture and detection Abs both recognize epitopes in A3-Nter (residues 17-165). ApoB-100 levels in VLDL fractions isolated from liver perfusates were measured using a commercial ELISA kit which specifically detects the C-terminal region of ApoB according to the manufacturer's instructions. Conditioned medium containing V5-tagged A3, either alone or co-expressed with A8, was immunoprecipitated with an anti-V5 antibody, and the precipitated proteins were quantified using a human A3 ELISA kit according to the manufacturer's instructions. The linearity of each assay was confirmed by serial dilution of the samples.

**Quantification of Circulating A3 by Selected Reaction Monitoring (SRM) Mass Spectrometry (MS)**

To improve detection of low-abundance plasma proteins, we first removed albumin from the samples using an albumin depletion kit. The samples were digested with trypsin and spiked in with 2 pmol of each heavy-isotope labeled peptide: SFVEQQDNSI**R** (aa166-167); TGIQEPSENSLSS**K** (aa206-219). Lys was labelled by $^{13}C_6$, $^{15}N_2$, and Arg was labeled by $^{13}C_6$, $^{15}N_4$. Peptides were selected based on their specificity and digestion efficiency. All peptides were synthesized by Life Technologies Corporation. Mass spectrometry was conducted in UTSW Proteomic Core.

**EL Inactivation by $A3_3$ and $A3_2A8$**

Structural models of $A3_2A8$ and $A3_3$ bound to EL or LPL were generated using the AlphaFold Server[58,59] using sequences corresponding to the secreted forms of human proteins: A3-FL (residues 17-460), A3-Nter (residues 17-221), A8 (residues 22-198), EL (residues 21-500) and LPL (residues 28-475). AlphaFold confidence metrics (ipTM/pTM) were used to assess model quality at the complex level. Because ipTM reflects confidence averaged across entire chains and the lipase-ANGPTL interaction interface is relatively small, we also generated "trimmed" models lacking low-confidence or disordered regions, EL (residues 46-485), A3 (35-152), and A8 (17-143). We refer to inhibitor-complex structures lacking disordered regions as "hinge" models because they are trimmed at a natural break, or hinge, in the N-terminal helix, allowing the remaining coiled-coil segment to bend at multiple angles (Table S5). Consistent with prior observations[59,60], the trimmed hinge models yielded the highest confidence metrics when complexed with EL, however, higher ipTM scores do not necessarily reflect confidence in the interaction interface itself. To better assess the EL interaction in both $A3_3$ and $A3_2A8$ complexes, we therefore focused on local interface quality, as reflected by AlphaFold3 inter-residue contact probabilities. Inter-residue contacts were defined as residue pairs with nonzero contact probabilities between EL and the primary interacting helix of the inhibitor complex, specifically, the A8 helix in the A3-$FL_2$A8 or the corresponding residues in the A3 helix (for the A3-$FL_2$A3-Nter complex, see heatmaps in Figure 6C), which more directly reflects the predicted strength and specificity of $A3_3$ and $A3_2A8$ binding surfaces. The sequence ranges of $A3_3$ and $A3_2A8$ with EL and LPL and their TM scores are provided in Table S5.

**Table S1. Oligonucleotides Sequences Used for Genotyping and Sequencing.**

| **Genotyping:** | |
|---|---|
| *A3*$^{FL/FL}$ | F-5'-TCTGCACATGCTCAGCAAGCACTGTGCCCC-3' |
| | R-5'-AGACAACAAGCCAGGGTCCAATTAAATTAAC-3' |
| *A3*$^{Nter/Nter}$ | F-5'-TCTGCACATGCTCAGCAAGCACTGTGCCCC-3' |
| | R-5'- TGGCAACTAGAAGGCACAGTCGAGG-3' |
| *A3*$^{-/-}$ | lacZ-F-5'- CGTGGCCTGATTCATTCC -3' |
| | lacZ-R-5'- ATCCTCTGCATGGTCAGGTC -3' |
| *A8* | WT (834 bp) |
| | F-5'-GCCTGCTTAGCCATCAGTCT-3' |
| | R-5'- GAAGGTCACTGAGGCTGGAA-3' |
| | KO (635 bp) |
| | F-5'-GCCTGCTTAGCCATCAGTCT-3' |
| | R-5'-GAAGGTCACTGAGGCTGGAA-3' |
| *Ldlr*$^{-/-}$ | WT |
| | F-5'- ACC AGA CGT GCT CCC AGG ATG A -3' |
| | R-5'- CGC AGT GCT CCT CAT CGT ACT TGT C -3' |
| | KO |
| | F-5'- GAT TGG GAA GAC AAT AGC AGG CAT GC -3' |
| | R-5'- GGC AAG ATG GCT CAG CAA GCA AAG GC -3' |
| *Lipg*$^{-/-}$ | WT |
| | F-5'-TTT TCC TCT TTC AAA GTG CTC CTG-3' |
| | R-5'-CCT ACG GGA ACT CCC TGA ACT TTT G -3' |
| | KO |

| | |
|---|---|
| | F-5'-TCA TTC TCA GTA TTG TTT TGC C -3' |
| | R-5'-AAG CCA GTG CTC ACC TAG C -3' |
| **Quantitative PCR** | |
| A3-(136-205) | F-5’-GCACCTTCAGAGCCAAAATCA-3’ |
| | R-5’-GCAGGAGGCCATTCGCTAA-3’ |
| A3-(2240-2314) | F-5’-GCATTCCCTAACTGCATATACAACA |
| | R-5’-ACCAGTCTTTCTGAGCTGATAAAAGAA |
| A3-(527-629) | F’-AGCAAGACAACAGCATAAGAGAACTC |
| | R’-CTGAGCTGCTTTTCTATTTCTTTTATCTG |
| A8-(434-547) | F-5’-CCCACCAAGAATTTGAGACCTT-3’ |
| | R-5’-ACTGTTGCTGCTCTGCCATCT-3’ |
| GAPDH-(230-231) | F-5’-TGTGTCCGTCGTGGATCTGA-3’ |
| | R-5’-GGTGCCTCTGGAGATTTTCG-3’ |
| 18S | F-5’-AGGGTTCGATTCCGGAGAGG-3’ |
| | R-5’-CAACTTTAATATACGCTATTGG-3’ |

177

**Table S2. Offspring of Heterozygous Matings of *A3^FL/FL^* mice**

| | Males (n=77) | | | | Females (n=69) | | | | | Total (n=146) | | | |
|---|---|---|---|---|---|---|---|---|---|---|---|---|---|
| | Observed | | Expected | | Observed | | Expected | | | Observed | | Expected | |
| | # | % | # | % | # | % | # | % | | # | % | # | % |
| **(+/+)** | 13 | 16.8 | 19.2 | 25 | 17 | 24.6 | 17.2 | 25 | **Males** | 77 | 55 | 73 | 50 |
| **(+/-)** | 44 | 57.1 | 38.5 | 50 | 34 | 49.3 | 34.5 | 50 | **Females** | 69 | 45 | 73 | 50 |
| **(-/-)** | 20 | 26.1 | 19.2 | 25 | 18 | 26.1 | 17.2 | 25 | | | | | |

| Breedings | | | Pups | | |
|---|---|---|---|---|---|
| Male | Female | No. of Litter | Pups per litter | Males | Females |
| (+/-) | (+/-) | 33 | 4.4 | 77 | 69 |
| (-/-) | (-/-) | 51 | 3.6 | 96 | 86 |
| (+/+) | (+/+) | 37 | 4.3 | 90 | 71 |

**Table S3. Offspring of Heterozygous Matings of *A3*$^{Nter/Nter}$ mice**

| | Males (n=39) | | | | Females (n=60) | | | | | Total (n=99) | | | |
|---|---|---|---|---|---|---|---|---|---|---|---|---|---|
| | Observed | | Expected | | Observed | | Expected | | | Observed | | Expected | |
| | # | % | # | % | # | % | # | % | | # | % | # | % |
| **(+/+)** | 10 | 25.6 | 9.7 | 25 | 19 | 31.6 | 15 | 25 | **Males** | 39 | 39.3 | 49.5 | 50 |
| **(+/-)** | 18 | 46.2 | 19.5 | 50 | 25 | 41.6 | 30 | 50 | **Females** | 60 | 60.6 | 49.5 | 50 |
| **(-/-)** | 11 | 28.2 | 9.7 | 25 | 16 | 26.7 | 15 | 25 | | | | | |

| Breedings | | | Pups | | |
|---|---|---|---|---|---|
| Male | Female | No. of Litter | Pups per litter | Males | Females |
| (+/-) | (+/-) | 19 | 5.2 | 39 | 60 |

**Table S4. Offspring of Heterozygous Matings of *A8*$^{-/-}$ mice**

| | Males (n=79) | | | | Females (n=62) | | | | | Total (n=141) | | | |
|---|---|---|---|---|---|---|---|---|---|---|---|---|---|
| | Observed | | Expected | | Observed | | Expected | | | Observed | | Expected | |
| | # | % | # | % | # | % | # | % | | # | % | # | % |
| **(+/+)** | 21 | 26.6 | 19.8 | 25 | 19 | 30.6 | 15.5 | 25 | **Males** | 79 | 56 | 70.5 | 50 |
| **(+/-)** | 39 | 49.3 | 39.5 | 50 | 31 | 50 | 31 | 50 | **Females** | 62 | 44 | 70.5 | 50 |
| **(-/-)** | 19 | 24.1 | 19.8 | 25 | 12 | 19.4 | 15.5 | 25 | | | | | |

| Breedings | | | Pups | | |
|---|---|---|---|---|---|
| Male | Female | No. of Litter | Pups per litter | Males | Females |
| (+/-) | (+/-) | 34 | 4.2 | 79 | 62 |

**Table S5. AlphaFold Structural Model Confidence**

| Enzyme | EL/LPL Residues | Inhibitor Complex | A3 Residues | A8 Residues | ipTM | pTM |
|---|---|---|---|---|---|---|
| NA | NA | A3-FL$_3$ | 17-460 | NA | 0.28 | 0.29 |
| NA | NA | A3-FL$_2$A3-Nter | 17-460; 17-221 | NA | 0.37 | 0.35 |
| NA | NA | A3-Nter$_3$* | 17-221 | NA | 0.54* | 0.54 |
| NA | NA | A3-Nter$_2$A8* | 17-221 | 22-198 | 0.52* | 0.52 |
| NA | NA | A3-FL$_2$A8 | 17-460 | 22-198 | 0.39 | 0.39 |
| EL | 21-500 | A3-FL$_3$ | 17-460 | NA | 0.27 | 0.35 |
| EL | 21-500 | A3-FL$_2$A3-Nter* | 17-460; 17-221 | NA | 0.37* | 0.45 |
| EL | 21-500 | A3-FL$_2$A8* | 17-460 | 22-198 | 0.42* | 0.46 |
| EL | 21-500 | A3-FLA3-NterA8 | 17-460; 17-221 | 22-198 | 0.46 | 0.51 |
| EL | 21-500 | A3-FLA3-Nter$_2$ | 17-460; 17-221 | NA | 0.42 | 0.47 |
| EL | 46-485 | A3A8 hinge | 35-152 | 27-143 | 0.49 | 0.68 |
| EL | 46-485 | A3 hinge | 35-152 | NA | 0.34 | 0.56 |
| LPL | 28-475 | A3-FL$_3$ | 17-460 | NA | 0.28 | 0.36 |
| LPL | 28-475 | A3-FL$_2$A3-Nter* | 17-460; 17-221 | NA | 0.32* | 0.39 |
| LPL | 28-475 | A3-FL$_2$A8* | 17-460 | 22-198 | 0.39* | 0.45 |

*Complex structure models were generated by substituting A8 with A3-Nter, which is similar in size and domain content.

**Table S6. Statistical analysis with exact P-values**

| | | |
|---|---|---|
| **Fig2C** | | |
| A3-FL: One-way ANOVA with Dunnett's multiple comparisons test | Summary | Adjusted P Value |
| +/+ vs. Full | + | 1.69E-08 |
| +/+ vs. Nter | ** | 6.02E-03 |
| +/+ vs. -/- | ** | 3.93E-03 |
| A3-Nter: One-way ANOVA with Dunnett's multiple comparisons test | Summary | Adjusted P Value |
| +/+ vs. Full | * | 1.03E-02 |
| +/+ vs. Nter | ** | 1.30E-03 |
| +/+ vs. -/- | ** | 4.90E-03 |
| A8: One-way ANOVA with Dunnett's multiple comparisons test | Summary | Adjusted P Value |
| +/+ vs. Full | * | 4.69E-02 |
| +/+ vs. Nter | ns | 4.49E-01 |
| +/+ vs. -/- | ns | 1.08E-01 |
| **Fig3A** | | |
| One-way ANOVA with Tukey's multiple comparisons test | Summary | Adjusted P Value |
| +/+ vs. FL/FL | + | 3.59E-07 |
| +/+ vs. Nter/Nter | *** | 2.29E-04 |
| +/+ vs. -/- | + | 6.40E-11 |
| FL/FL vs. Nter/Nter | *** | 9.63E-04 |
| FL/FL vs. -/- | + | 1.29E-06 |
| Nter/Nter vs. -/- | + | 2.39E-09 |
| **Fig 3B** | | |
| One-way ANOVA- Fisher's LSD | | |
| +/+ vs FL/FL | ** | 1.15E-03 |
| +/+ vs Nter/Nter | * | 1.19E-02 |
| FL/FL vs FL/Nter | + | 7.19E-06 |
| Nter/Nter vs FL/Nter | + | 9.80E-05 |
| **Fig 3C** | | |
| One-way ANOVA with Tukey's multiple comparisons test | Summary | Adjusted P Value |
| +/+ vs. FL/FL | ns | 9.93E-01 |
| +/+ vs. Nter/Nter | * | 2.40E-02 |
| +/+ vs. -/- | + | 1.30E-05 |
| FL/FL vs. Nter/Nter | * | 1.42E-02 |
| FL/FL vs. -/- | + | 8.62E-06 |
| Nter/Nter vs. -/- | ** | 2.17E-03 |
| **Fig 3E** | | |
| One-way ANOVA- Fisher's LSD | | |

| | | |
|---|---|---|
| +/+: AAV-A8 vs GFP | + | 1.01E-07 |
| FL: AAV-A8 vs GFP | *** | 5.22E-04 |
| Nter: AAV-A8 vs GFP | + | 2.15E-06 |
| +/+-AAV-A8 vs FL-AAV-A8 | + | 1.00E-05 |
| +/+-AAV-A8 vs Nter-AAV-A8 | ns | 7.60E-02 |
| FL-AAV-A8 vs Nter-AAV-A8 | *** | 8.46E-04 |
| **Fig 5B, left** | | |
| Nonparametric Mann-Whitney test (unpaired, two tail) | | |
| *Ldlr*$^{-/-}$ *vs Ldlr*$^{-/-}$*;A8*$^{-/-}$ | * | 2.80E-02 |
| **Fig 5C** | | |
| Unpaired two-tailed t test | | |
| Panel: left | * | 1.08E-02 |
| Panel: right | ** | 3.53E-03 |
| **Fig 5D_top panel** | | |
| One-way ANOVA Tukey's multiple comparisons test | | |
| *Ldlr*$^{-/-}$ *vs Ldlr*$^{-/-}$*;A8*$^{-/-}$ | *** | 7.11E-04 |
| *Ldlr*$^{-/-}$*;A8*$^{-/-}$ *vs Ldlr*$^{-/-}$*;A8*$^{-/-}$*;Lipg*$^{-/-}$ | ns | 7.15E-01 |
| *Ldlr*$^{-/-}$*vs Ldlr*$^{-/-}$*;A8*$^{-/-}$*;Lipg*$^{-/-}$ | *** | 7.82E-04 |
| **Fig 5D_bottom panel** | | |
| One-way ANOVA Tukey's multiple comparisons test | | |
| *Ldlr*$^{-/-}$ *vs Ldlr*$^{-/-}$*;A8*$^{-/-}$ | * | 2.62E-02 |
| *Ldlr*$^{-/-}$*;A8*$^{-/-}$ *vs Ldlr*$^{-/-}$*;A8*$^{-/-}$*;Lipg*$^{-/-}$ | *** | 3.70E-04 |
| *Ldlr*$^{-/-}$*vs Ldlr*$^{-/-}$*;A8*$^{-/-}$*;Lipg*$^{-/-}$ | * | 2.10E-02 |
| **Fig 5F** | | |
| One-way ANOVA- Fisher's LSD | | |
| column 1 VS 3 | ** | 8.17E-03 |
| column 1 VS 5 | + | 1.90E-05 |
| column 3 VS 4 | + | 2.80E-05 |
| **Fig 5G** | | |
| One-way ANOVA- Fisher's LSD | Summary | Individual P Value |
| *Ldlr*$^{-/-}$ vs. *Ldlr*$^{-/-}$*;A3*$^{-/-}$ | + | 5.19E-06 |
| *Ldlr*$^{-/-}$ vs. *Ldlr*$^{-/-}$*;A8*$^{-/-}$ | *** | 1.37E-04 |
| *Ldlr*$^{-/-}$ vs. *Ldlr*$^{-/-;}$*A3*$^{-/-}$*;A8*$^{-/-}$ | + | 8.32E-06 |
| *Ldlr*$^{-/-}$*;A3*$^{-/-}$ vs. *Ldlr*$^{-/-}$*;A8*$^{-/-}$ | * | 1.48E-02 |
| *Ldlr*$^{-/-}$*;A3*$^{-/-}$ vs. *Ldlr*$^{-/-;}$*A3*$^{-/-}$*;A8*$^{-/-}$ | ns | 7.08E-01 |
| *Ldlr*$^{-/-}$*;A8*$^{-/-}$ vs. *Ldlr*$^{-/-;}$*A3*$^{-/-}$*;A8*$^{-/-}$ | * | 3.13E-02 |
| **Fig 6B** | | |
| Paired two-tail t test | | |
| 12.5 ng, A3: A3+A8 | ** | 3.00E-03 |
| 25 ng, A3: A3+A9 | * | 3.39E-02 |

| | | |
|---|---|---|
| **Fig 6D** | | |
| **Left panel** | | |
| One-way ANOVA- Fisher's LSD | Summary | Individual P Value |
| Column 1 vs. 2 | ** | 9.10E-03 |
| Column 1 vs. 4 | * | 4.67E-02 |
| Column 2 vs. 3 | ** | 1.10E-03 |
| Column 2 vs. 4 | ns | 2.83E-01 |
| Column 3 vs. 4 | ** | 5.70E-03 |
| **Right panel** | | |
| One-way ANOVA- Fisher's LSD | Summary | Individual P Value |
| Column 2 vs. 3 | *** | 1.00E-04 |
| Column 2 vs. 4 | ** | 1.46E-03 |
| Column 3 vs. 4 | ** | 2.60E-03 |
| **Fig S2B** | | |
| left, One-way ANOVA- Fisher's LSD | Summary | Individual P Value |
| +/+ vs. Nter | + | 3.15E-07 |
| FL vs. Nter | + | 5.82E-07 |
| Right, One-way ANOVA- Fisher's LSD | Summary | Individual P Value |
| +/+ vs. Nter | + | 1.68E-09 |
| FL vs. Nter | + | 6.96E-10 |
| **Fig S2C** | | |
| One-way ANOVA- Fisher's LSD | Summary | Individual P Value |
| +/+ vs. FL/FL | + | 2.53E-08 |
| +/+ vs. Nter/Nter | * | 4.93E-02 |
| FL/FL vs. Nter/Nter | + | 5.97E-09 |
| **Fig S2E** | | |
| Left: One-way ANOVA- Fisher's LSD | Summary | Individual P Value |
| WT vs. A3-Full | *** | 9.43E-04 |
| WT vs. A3-N | * | 1.75E-02 |
| A3-Full vs. A3-N | + | 4.50E-05 |
| Right: Unpaired two-tailed t test | | |
| A3-Full vs. A3-N | + | 1.00E-05 |
| **Fig S2F** | | |
| One-way ANOVA- Fisher's LSD | Summary | Individual P Value |
| +/+ vs. FL | ** | 1.08E-03 |
| FL vs. Nter | *** | 7.00E-04 |
| **Fig S3B** | | |
| Left: One-way ANOVA- Fisher's LSD | Summary | Individual P Value |
| WT vs. FL/- | * | 1.28E-02 |
| WT vs. FL/FL | **** | 4.69E-09 |
| WT vs. FL/Nter | ** | 1.39E-03 |

| | | |
|---|---|---|
| FL/FL vs. FL/Nter | **** | 1.32E-06 |
| Right: One-way ANOVA- Fisher's LSD | Summary | Individual P Value |
| WT vs. Nter/- | ** | 1.60E-03 |
| Nter/- vs. Nter/Nter | ** | 5.70E-03 |
| Nter/- vs. FL/Nter | ** | 2.50E-03 |
| **Fig S5** | | |
| Nonparametric Mann-Whitney test (unpaired, two tail) | | |
| left: Ldlr$^{-/-}$ vs. Ldlr$^{-/-}$;A8$^{-/-}$ | ns | 5.00E-01 |
| Right: Ldlr$^{-/-}$ vs. Ldlr$^{-/-}$;A8$^{-/-}$ | * | 2.32E-02 |
| **Fig S6B** | | |
| One-way ANOVA- Fisher's LSD | | |
| Column 1 vs. 2 | + | 8.60E-09 |
| Column 3 vs. 4 | + | 2.88E-10 |
| **Fig S6C** | | |
| One-way ANOVA- Fisher's LSD | Summary | Individual P Value |
| *Ldlr*$^{-/-}$ vs. *Ldlr*$^{-/-}$;*A3*$^{-/-}$ | + | 1.20E-05 |
| *Ldlr*$^{-/-}$ vs. *Ldlr*$^{-/-}$;*A8*$^{-/-}$ | + | 1.00E-05 |
| *Ldlr*$^{-/-}$ vs. *Ldlr*$^{-/-;}$*A3*$^{-/-}$;*A8*$^{-/-}$ | + | 1.50E-05 |
| **Fig S6D** | | |
| One-way ANOVA- Fisher's LSD | Summary | Adjusted P Value |
| **Triglyceride** | | |
| WT vs. FL/FL | *** | 3.51E-04 |
| WT vs. Nter/Nter | ** | 2.58E-03 |
| FL/FL vs. Nter/Nter | ns | 8.71E-02 |
| **Cholesterol** | | |
| WT vs. FL/FL | ns | 2.80E-01 |
| WT vs. Nter/Nter | ** | 3.96E-03 |
| FL/FL vs. Nter/Nter | ns | 1.28E-01 |

### Figure S1

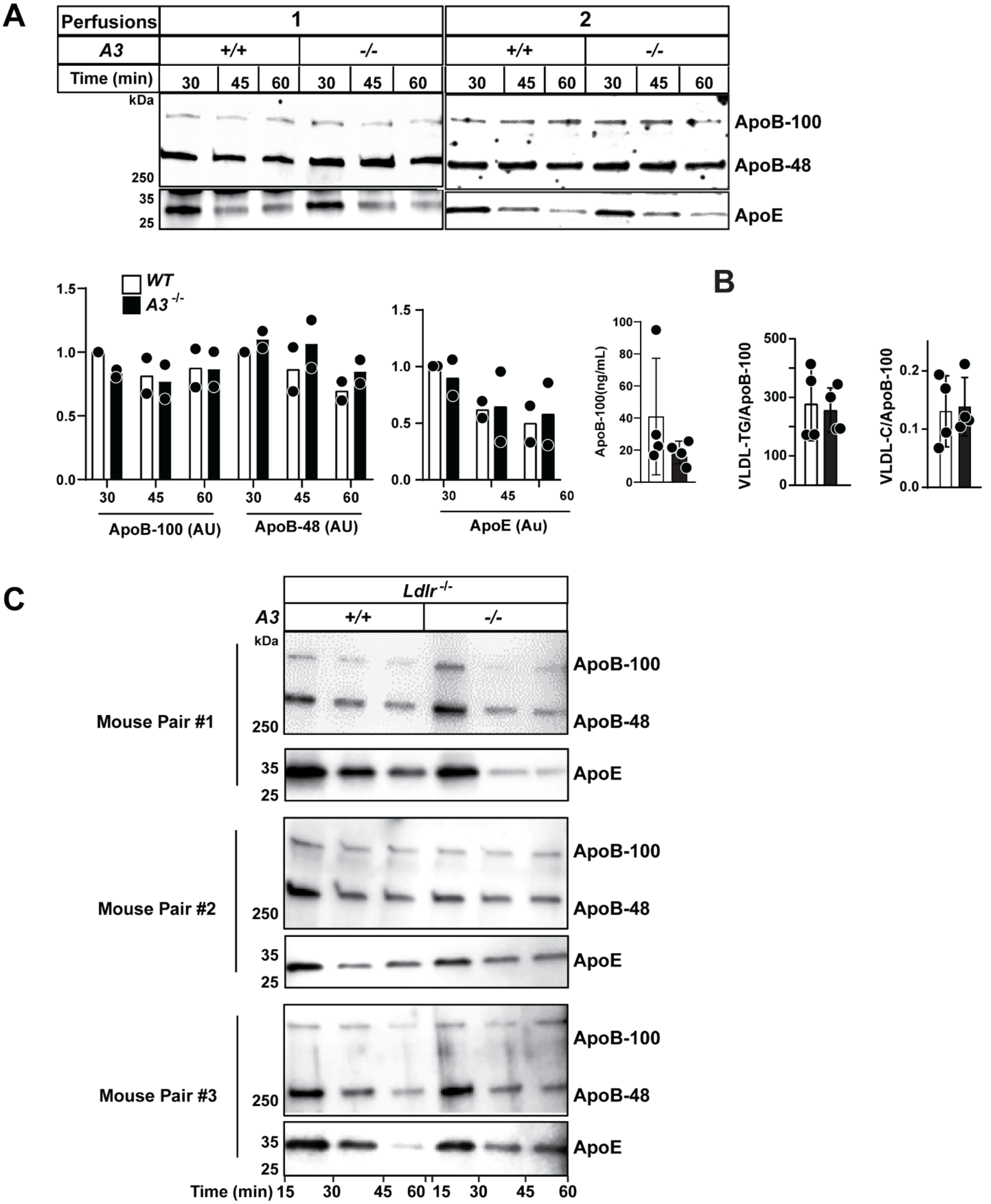

A
Perfusions
1
2
A3
+/+
-/-
Time (min)
30
45
60
kDa
250
35
25
ApoB-100
ApoB-48
ApoE
WT
A3 -/-
ApoB-100 (AU)
ApoB-48 (AU)
ApoE (Au)
ApoB-100(ng/mL)
B
VLDL-TG/ApoB-100
VLDL-C/ApoB-100
C
Ldlr -/-
Mouse Pair #1
Mouse Pair #2
Mouse Pair #3
Time (min) 15 30 45 60 15 30 45 60

**Figure S1. Liver perfusion in WT and *A3*$^{-/-}$ mice without heparin treatment. A,** Immunoblot analysis of ApoB and ApoE in isolated VLDL was performed as described in the Supplementary Methods (male, n=4/group, 10-12 wks, ad-lib fed). ApoB-100 levels were quantified by LI-COR(vales are normalized to the +/+ group at 30 min timepoint) and ELISA. **B,** Lipid content in VLDL measured enzymatically and normalized to ApoB-100 levels. **C**. Immunoblot of VLDL-ApoB and -ApoE in *Ldlr*$^{-/-}$;*A3*$^{-/-}$ and *Ldlr*$^{-/-}$ littermates (n=3/group, age matched 8-19 wks) after 2 wks on a high-sucrose diet. Each perfusion experiment included one WT and one *A3*$^{-/-}$ mouse. Mann-Whitney test was performed between WT and KO groups (A, right and B) (n=4), no significant differences were observed ($p>0.05$ for all comparisons). Four independent experiments were performed with similar results.

### Figure S2

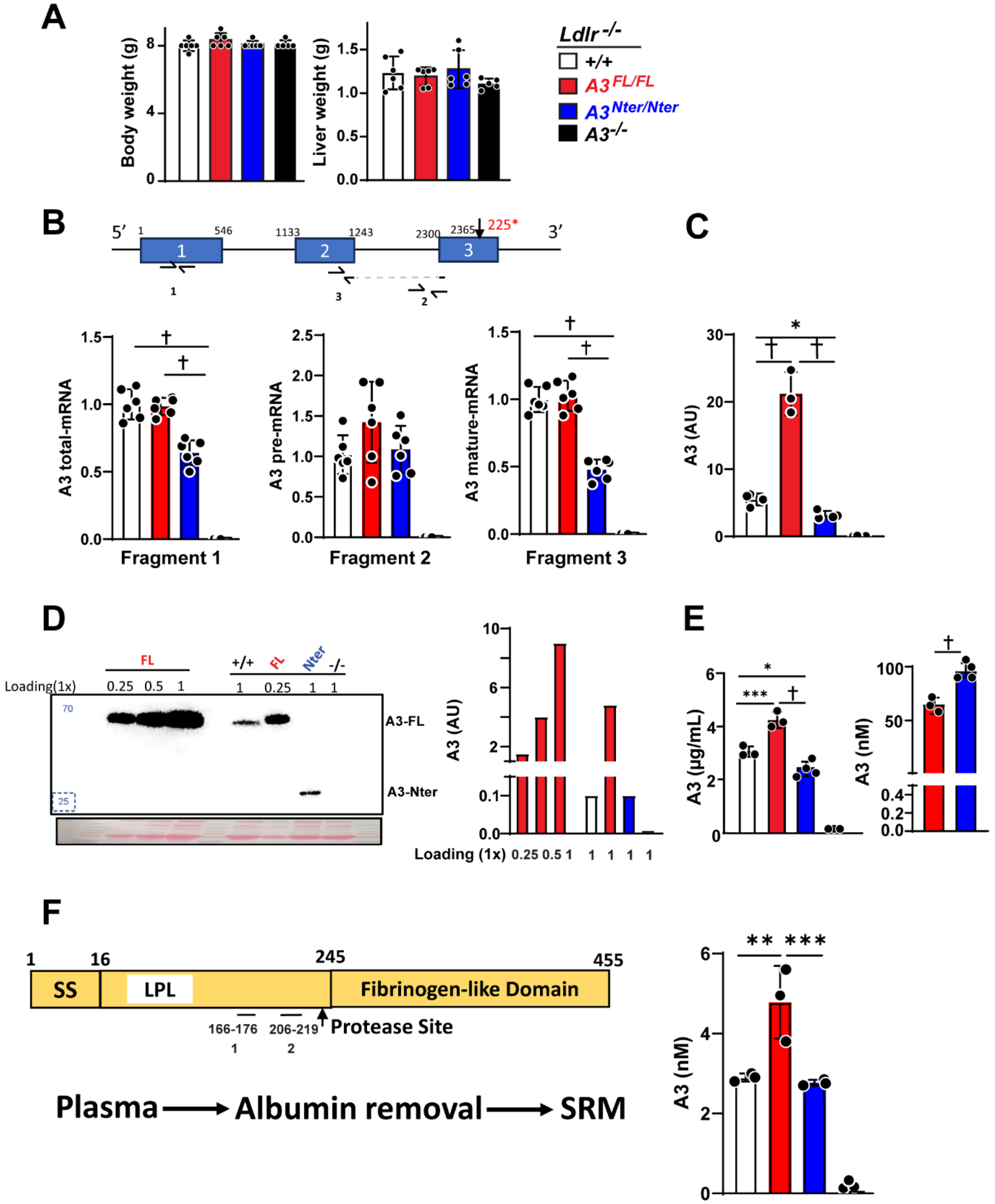

A
B
C
D
E
F
Body weight (g)
Liver weight (g)
Ldlr -/-
+/+
A3 FL/FL
A3 Nter/Nter
A3 -/-
5′
3′
225*
A3 total-mRNA
A3 pre-mRNA
A3 mature-mRNA
Fragment 1
Fragment 2
Fragment 3
A3 (AU)
Loading(1x)
A3-FL
A3-Nter
Loading (1x)
A3 (µg/mL)
A3 (nM)
SS
LPL
Fibrinogen-like Domain
Protease Site
166-176
206-219
Plasma
Albumin removal
SRM

**Figure S2. Effect of A3 cleavage on body weight, liver weight, mRNA and A3 protein levels.** **A,** Body weights (left) and liver weights (right) of mice expressing A3 isoforms (male, n=6, 6, 6, 5/group, 11-16 wks of age, refed 4 h). **B**, Messenger RNA (mRNA) levels of A3 in indicated mouse strains. Top, Diagram of A3 with location of primers used for qRT-PCR. Bottom, qRT-PCR analysis of A3 mRNA levels in livers of refed male nice (8-12 wks old, n=6, 6, 6 and 5/group). Values are normalized to the +/+ group. **C**. Band intensities of A3-FL and A3-Nter were quantified by LI-COR in the indicated refed mice (males, 8-12 wks, n=4, 3, 4, 4/group). In the $A3^{FL/FL}$ mice, we included the signal at the expected molecular mass of the A3-Nter band; likewise, in the $A3^{Nter/Nter}$ mice, we included the signal at the expected molecular mass of A3-FL. Total A3 was calculated as the sum of A3-FL and A3-Nter in all mouse models. **D.** Immunoblot of plasma using an anti-A3-Nter mAb. Immunoblot analysis was performed using 1 μL of plasma from the WT and A3-Nter mice and 0.25 μL from the A3-FL mice. Serial dilutions of plasma from the A3-FL were made using saline to establish the range in which the levels of detection were linear (left). **E**. Quantification of A3 in plasma by ELISA. Plasma was serially diluted to define the linear range of detection. Molarity of A3 in mice expressing A3-FL or A3-Nter was calculated from the mass measured by ELISA and the molecular weight of the peptides (65 and 27 kDa, respectively). Males, 7-9 wks, n=3, 3, 4, 3/group, refed 4 h. **F.** Quantification of A3 levels in plasma by SRM mass spectrometry. Left: Two peptides labeled with $^{13}C$ and $^{15}N$, were used as standards to quantify the levels of A3 protein. Right: Circulating A3 levels were measured in plasma from refed male mice (8-12 wks, 3/group) after removal of albumin. The mean levels of A3 were determined by taking the mean of the three peptide measurements. Mean values were compared using nonparametric Kruskal-Wallis test followed by Dunn's procedure (A), parametric unpaired two-tailed student-t test (E, right) or one way ANOVA-Fisher's LSD test (B, E, F) in GraphPad Prism. All experiments were repeated 3 times with similar results. Mean ± S.D. * $p < 0.05$ ** $p < 0.01$ *** $p < 0.001$ + $p < 0.0001$. Statistical values were not added to the graphs when no significant differences were found. When differences were statistically significant, the exact $p$ values were provided in Supplementary Table 6**.**

### Figure S3

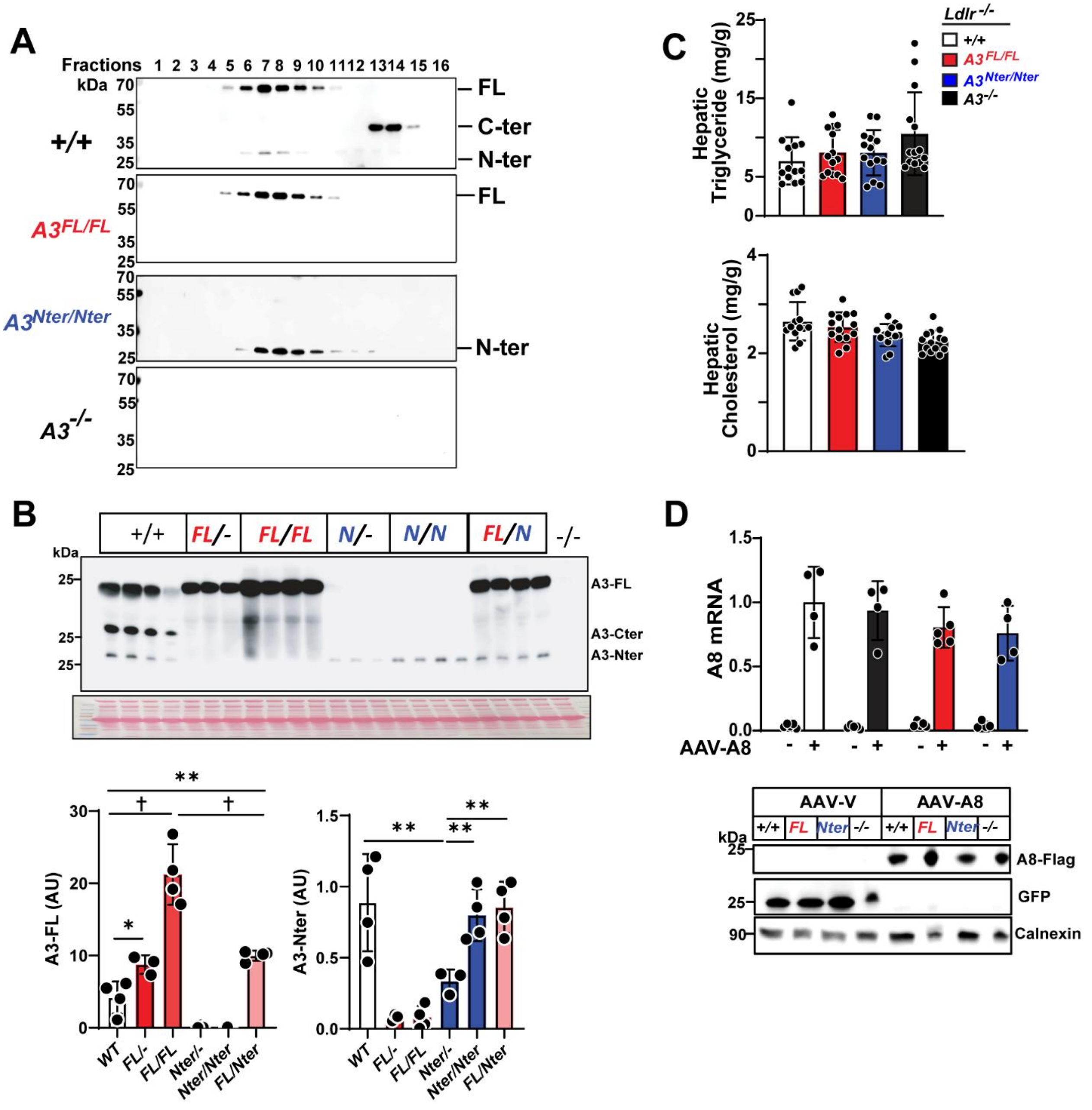


**Figure S3. Effect of A3 cleavage on A3 hepatic lipid levels. A.** Immunoblot of A3 in plasma fractions (12.5 µL per lane) separated by Superdex 200 size-exclusion chromatography (male, n=3, 3, 4, and 4/group, 8-12 wks, refed 4 h). **B**. Immunoblot analysis of plasma A3 in indicated mouse models. Assays were performed using plasma from refed age-matched male mice (n=4, 3, 4, 3, 4 and 4/group, 6-13 wks). Plasma was diluted 1:10 in saline and a total of 10 µl was loaded per lane.

Protein levels were quantified by LI-COR. **C**, Hepatic TG and cholesterol levels in 4 strains of mice. Data from 3 independent experiments were pooled (male, n=13, 14, 14, and 14/group, age matched 9-16 wks of age, refed 4 h). **D**, Hepatic *A8* mRNA and protein levels following A8 overexpression as described in Supplementary Methods (male, n=4, 4, 5, 4, 5, 5, 3, and 4/group, 8-12 wks, fast 16 h). Values are normalized to column 2 (+/+-AAV-A8 group). Mean values were compared using one-way ANOVA with Tukey' test (C) or one-way ANOVA with Fisher's LSD test (B, D). Mean ± S.D. * $p < 0.05$ ** $p < 0.01$ + $p < 0.0001$. Statistical values were not added to graphs when no significant difference was observed (p>0.05). When differences were statistically significant, the exact *p* values were provided in Supplementary Table 6. All experiments were repeated 3 times with similar results.

### Figure S4

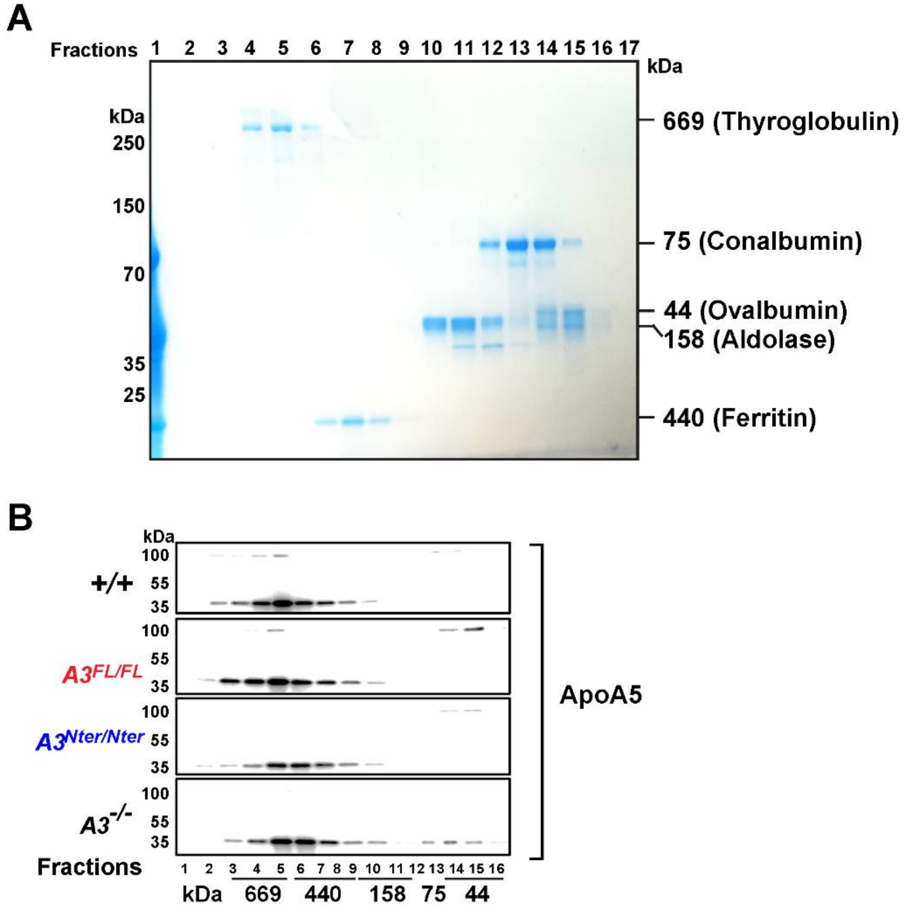


**Figure S4. A3-FL and A3-Nter co-elute with ApoA5 following size-exclusion chromatography. A**, Coomassie blue staining of molecular weight standards separated by a size exclusion column Superdex 200. A mixture of molecular weight standard makers was injected onto the column and fractionated using the same protocol as used for plasma samples. Odd-numbered fractions were collected, subjected to SDS-PAGE, and then stained with Coomassie blue. **B**, Immunoblot analysis of ApoA5 in plasma fractions separated by the size exclusion column Superdex 200 (male, n=6, 4, 6, and 6/group, 8-12 wks, refed 4 h). Experiments were repeated 3 times with similar results.

### Figure S5

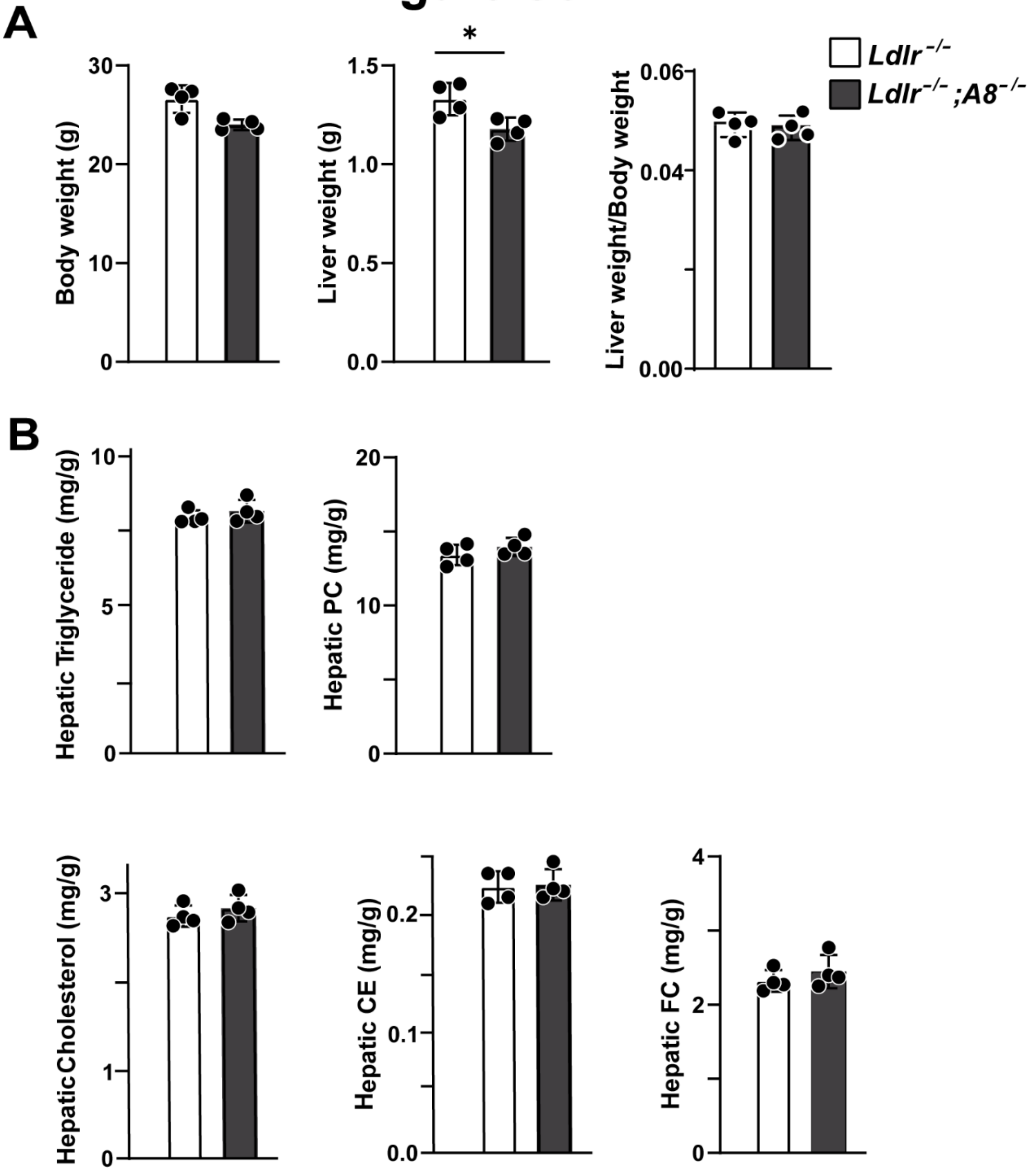


**Figure S5. Effects of A8 inactivation on body, liver weight and hepatic lipid levels in *Ldlr*$^{-/-}$ mice. A,** Body weight (left) and liver weight (right) in *Ldlr*$^{-/-}$ and *Ldlr*$^{-/-}$;A8$^{-/-}$ mice (male, n=4/group, 11-13 wks, refed 4 h). *$p$=0.03. **B**, Hepatic lipid levels measured in the same strains of mice. Data is shown as mean ± SD. Statistical significance was determined using non-parametric Mann-Whitney test (unpaired, two-tailed) in GraphPad Prism. Statistical values were not added to the graphs when no significant difference was observed ($p>0.05$). Experiments were repeated twice with similar results.

### Figure S6

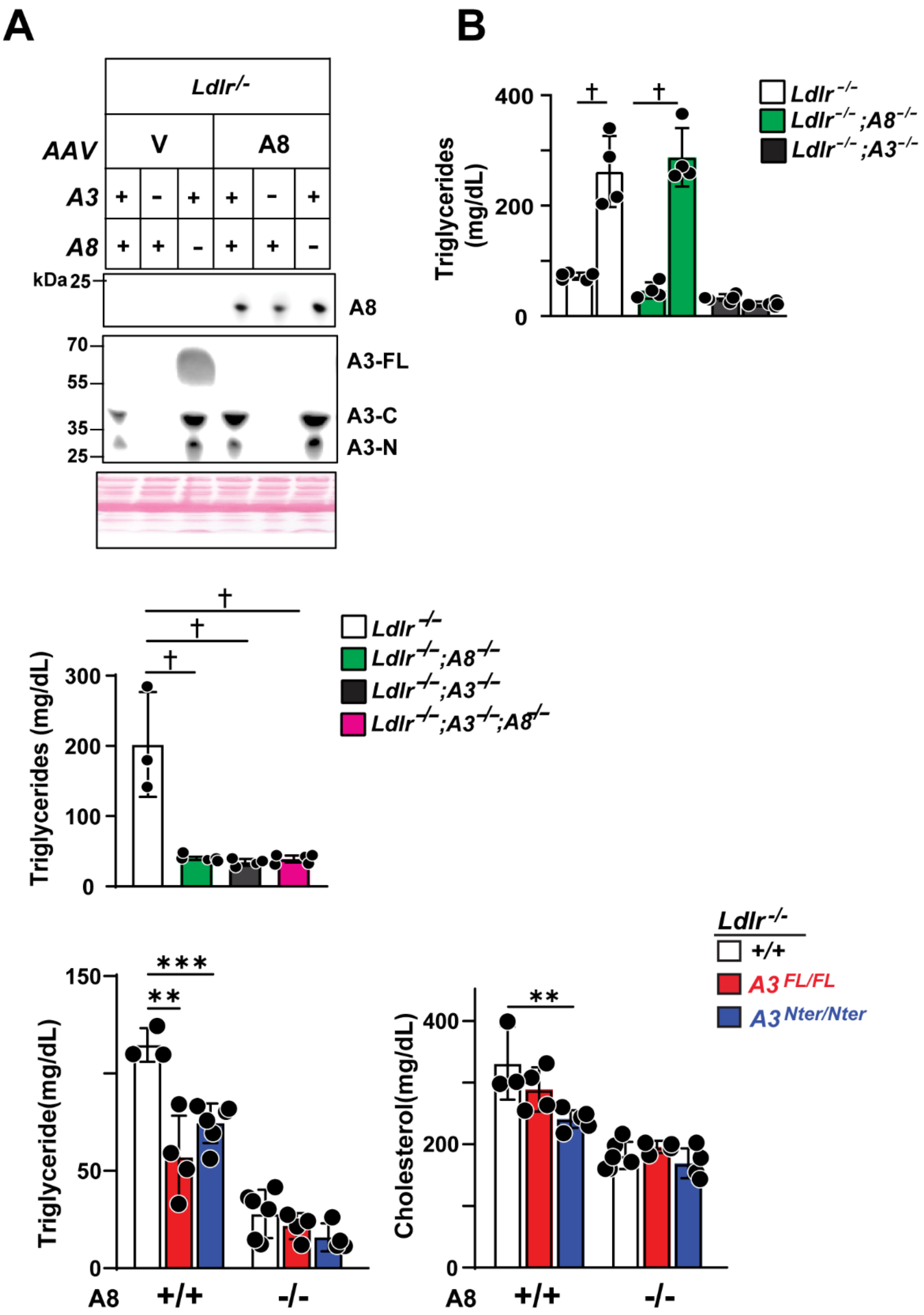


261

**Figure S6. Both A3 and A8 are necessary to increase plasma levels of TG and cholesterol. A,** Protein levels of circulating A8 and A3 and **B**, plasma TG levels following hepatic overexpression of A8 in *Ldlr*$^{-/-}$, *Ldlr*$^{-/-}$*;A8*$^{-/-}$, *Ldlr*$^{-/-}$*;A3*$^{-/-}$ mice, as described in Supplementary Methods and Figure 5F (male, n=5, 4, 4, 4, 5, and 5/group, 8-12 wks, fast 16 h). A FLAG antibody was used to detect A8, and AF136 pAb was used to detect A3 (10 μL per lane). **C**, Plasma TG levels in mice lacking both A3 and A8 (male, n=3, 5, 4, and 4/group, 8-11 wks, refed 4 h). **D,** Plasma TG and cholesterol levels in *Ldlr*$^{-/-}$, *Ldlr*$^{-/-}$*;A3*$^{FL/FL}$, *Ldlr*$^{-/-}$*;A3*$^{Nter/Nter}$ mice expressing or lacking *A8* (male, n=3, 4, 6, 6, 4, and 4/group, age matched 6-14 wks, refed 4 h). Mean values were compared using one -way ANOVA-Fisher LSD (B-D) in GraphPad Prism. Data are shown as mean ± SD. **p*$<$ 0.05, ***p*$<$ 0.01, ****p*$<$0.001, †*p*$<$0.0001. Exact *p* values were provided in Supplementary Table 6. Experiments were repeated twice with similar results.

### Figure S7

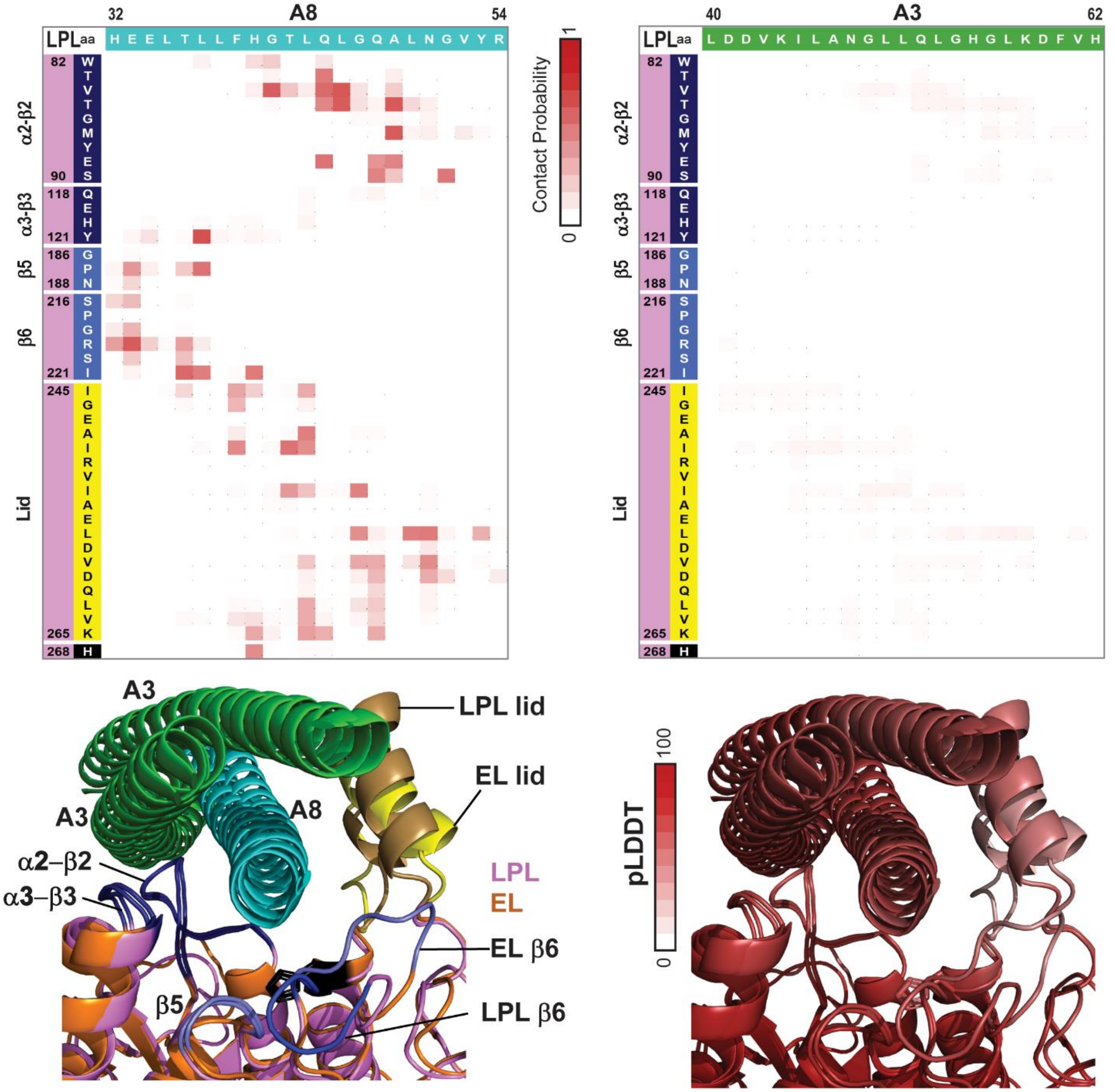


Figure S7. **Structural models of LPL: $A3_3$ and LPL: $A3_2A8$**. Top: Contact probabilities for residues in LPL with A8 (left) and A3 (right). Color scale bar: 0-1, units: contact probability. Bottom: superposition of EL and LPL complexes with $A3_2A8$ colored by structure element (left)

and complexes colored by pLDDT confidence values for AF models in white-red scale from 0 to 100 (right).
